# Guiding functional near-infrared spectroscopy optode-layout design using individual (f)MRI data: Effects on signal quality and sensitivity

**DOI:** 10.1101/2020.09.27.315390

**Authors:** A. Benitez-Andonegui, M. Lührs, L. Nagels-Coune, D. Ivanov, R. Goebel, B. Sorger

**Affiliations:** Maastricht Brain Imaging Center, Department Cognitive Neuroscience, Maastricht University, Oxfordlaan 55, Maastricht, the Netherlands, 6229 EV; Laboratory for Cognitive Robotics and Complex Self-Organizing Systems, Department of Data Science and Knowledge Engineering, Paul-Henri Spaaklaan 1, Maastricht University, Maastricht, the Netherlands, 6229 EN; Research Department, Brain Innovation B.V., Oxfordlaan 55, Maastricht, the Netherlands, 6229 EV

**Keywords:** functional near-infrared spectroscopy, functional magnetic resonance imaging, optode layout design, mental imagery

## Abstract

Designing optode layouts is an essential step for functional near-infrared spectroscopy (fNIRS) experiments as the quality of the measured signal and the sensitivity to cortical regions-of-interest depend on how optodes are arranged on the scalp. This becomes particularly relevant for fNIRS-based brain-computer interfaces (BCIs), where developing robust systems with few optodes is crucial for clinical applications. Available resources often dictate the approach researchers use for optode-layout design. Here we compared four approaches that incrementally incorporated subject-specific magnetic resonance imaging (MRI) information while participants performed mental-calculation, mental-rotation and inner-speech tasks. The literature-based approach (LIT) used a literature review to guide the optode layout design. The probabilistic approach (PROB), employed individual anatomical data and probabilistic maps of functional MRI (fMRI)-activation from an independent dataset. The individual fMRI (iFMRI) approach used individual anatomical and fMRI data, and the fourth approach used individual anatomical, functional and vascular information of the same subject (fVASC). The four approaches resulted in different optode layouts and the more informed approaches outperformed the minimally informed approach (LIT) in terms of signal quality and sensitivity. Further, PROB, iFMRI and fVASC approaches resulted in a similar outcome. We conclude that additional individual MRI data leads to a better outcome, but that not all the modalities tested here are required to achieve a robust setup. Finally, we give preliminary advice to efficiently using resources for developing robust optode layouts for BCI and neurofeedback applications.

## 1 Introduction

Functional near-infrared spectroscopy (fNIRS) is a non-invasive, portable optical imaging method used to measure brain activity via hemodynamic responses involving increased oxygen consumption and cerebral blood flow ^1–3^. These physiological changes lead to local changes in the concentrations of oxy-(Δ[HbO]) and deoxy-hemoglobin (Δ[HbR]), which can be detected because near-infrared light is absorbed by hemoglobin located in blood vessels ^3, 4^.

When setting up an fNIRS experiment, optical sensors (‘optodes’) are placed on the scalp, which can be classified into sources (emitters) and detectors (receivers). Light emitted from a source is propagated through extracerebral and cerebral tissues up to a few centimeters, where some photons are scattered and absorbed before light reaches the detectors ^5^. The spatial resolution of fNIRS is therefore in the range of 5-10mm ^4^ depending on the way source-detector pairs (or ‘channels’) are arranged on the scalp ^6^. The distance between a source and detector pair, along with the anatomical tissues between them determines the depth of light penetration and the sensitivity to underlying cortex ^1^. Therefore, the quality of the fNIRS signal can differ dramatically between optode layouts.

This effect of optode layout is particularly relevant for applications requiring sparse optode layouts, such as brain-computer interfaces (BCIs). BCIs provide an alternative means of motor-independent communication for clinical populations suffering from severe motor disabilities ^7^ by enabling users to send commands via brain activity in the absence of motor output ^7, 8^. FNIRS is a promising choice for implementing BCIs due to its portability, safety and relatively low cost ^9, 10^. However, it remains a challenging undertaking to develop efficient, accurate and robust systems using the limited number of optodes required for fNIRS-BCI systems to remain portable and comfortable for clinical applications. Indeed, a number of fNIRS-based BCI studies using small optode layouts ^11–16^ have reported variability in the number of participants able to reach the minimum accuracy (70% in a two-class BCI) required for practical BCI use ^17^. This variability may originate from individual anatomical ^18, 19^ or functional differences ^15^ that affect fNIRS signal quality/sensitivity and therefore might be improved by designing optode layouts for individual users that account for such differences.

Researchers often define a region of interest (ROI) in line with their research question and design an optode layout in a grid-like fashion to target a specific brain area ^1^. The simplest and most common optode-layout design is to assign source and detector locations on the head to cover a given cortical ROI according to the standardized 10-20 electroencephalography (EEG) system or its extended versions ^20^. These locations can be related to the underlying assumed cortical structure ^21, 22^ or to the standard Montreal Neurological Institute (MNI) stereotactic coordinates ^23–26^. This procedure has proven effective for many applications but may be suboptimal for use in BCIs. In this study, we were interested in whether incorporating additional neuroimaging data such as anatomical or functional magnetic resonance imaging (MRI or fMRI) can improve optode-layout design for use in BCIs.

The selection of the ROIs in the procedure described above are commonly based on anatomically defined coordinates only. However, ROIs derived from functional neuroimaging techniques such as fMRI could increase the spatial specificity of ROI definition by accounting for individual local differences in elicited brain activity for a given task. Once an ROI is defined, the fNIRS community has developed several approaches to optimize optode-layout designs using light-sensitivity profiles ^1^. Light-sensitivity profiles are probabilistic models of photon absorption based on the tissues found between source and detector optodes ^27^. Software packages, toolboxes and pipelines compute these profiles using Monte Carlo simulations to optimize optode layouts ^1, 5, 27–30^, thus promising an increase on signal quality and sensitivity for BCI applications. However, light sensitivity profile models require anatomical head data, either from an MRI-derived atlas or from subject-specific MRI data. MRI atlases are an appealing option for computing profiles, as they do not require additional MRI measurements, which may be expensive, time-consuming or generally unavailable. That said, subject-specific MRI data better capture specific anatomical and vascular features and therefore could improve the robustness of fNIRS setups across individuals. Considering subject-specific vascular information may be particularly relevant as vascular structures are highly scattering and absorbing media ^31^ and can influence the estimates of light sensitivity profiles ^32^.

Naturally, available resources for collecting additional data must dictate the approach researchers use to design optode layouts. We therefore asked the following question: What is the potential gain of incorporating (anatomical, functional, vascular) MRI data when optimizing optode-layout designs for fNIRS-based BCIs? With this question in mind, we selected four approaches that incrementally incorporated the amount of individual information from the same participant to design subject-specific optode layouts. The first layout was the literature-based approach (hereinafter referred to as LIT), where optodes were selected based on a literature review. LIT represents the scenario where no additional individual MRI information is available. The second setup was the probabilistic approach (referred to as PROB), which employed individual anatomical data together with a probabilistic functional map derived from an independent dataset to inform optode placement. PROB illustrates a situation where individual fMRI data is not available, but subject-specific anatomical information and functional data from other individuals is accessible. The third setup was the individual fMRI approach, which used anatomical data and functional activation maps of the same individual (referred to as iFMRI). Finally, the fourth setup was the vascular approach, which used individual anatomical, functional and vascular information of the same subject (referred to as fVASC).

We assessed whether different approaches resulted in distinct optode layouts and assessed whether the quality of the fNIRS signal and the detected task-related activation (fNIRS sensitivity) differed across optode layouts. Participants were asked to perform three mental-imagery tasks commonly used for hemodynamic BCIs, see Table S3: mental-calculation, mental-rotation and inner-speech. We designed approach-specific optode layouts using Monte Carlo simulations and an algorithmic procedure that used two main constraints: 1) the inter-optode distance did not exceed the 25-40mm range in order to provide a reasonable signal-to-noise ratio ^33^ and 2) the optode layout for each approach consisted of two channels that shared a common source. Importantly, the second constraint allowed us to compare the four approaches within the same functional fNIRS run. We hypothesized that each approach would lead to different optode-layout designs and that the signal-to-noise ratio of resulting fNIRS signal would improve with more individualized approaches. Our results show that the four approaches indeed result in different optode layouts and that the more individualized approaches (PROB, iFMRI, and fVASC) outperform the minimally informed approach (LIT) in terms of fNIRS signal quality and sensitivity. Further, we find that PROB, iFMRI, and fVASC approaches produce similar signal quality and sensitivity. Finally, we give preliminary recommendations to help researchers efficiently use resources for developing robust and convenient optode layouts for fNIRS-BCIs.

## 2 Materials and Methods

This experiment consisted of three separate sessions that took place in the following order: one f/MRI session, a neuronavigation session and an fNIRS session. The first two sessions aimed at gathering necessary information for designing optode layouts, while the fNIRS session aimed at acquiring data to assess/compare the designed optode layouts (see Fig. 1).

**Fig 1.**
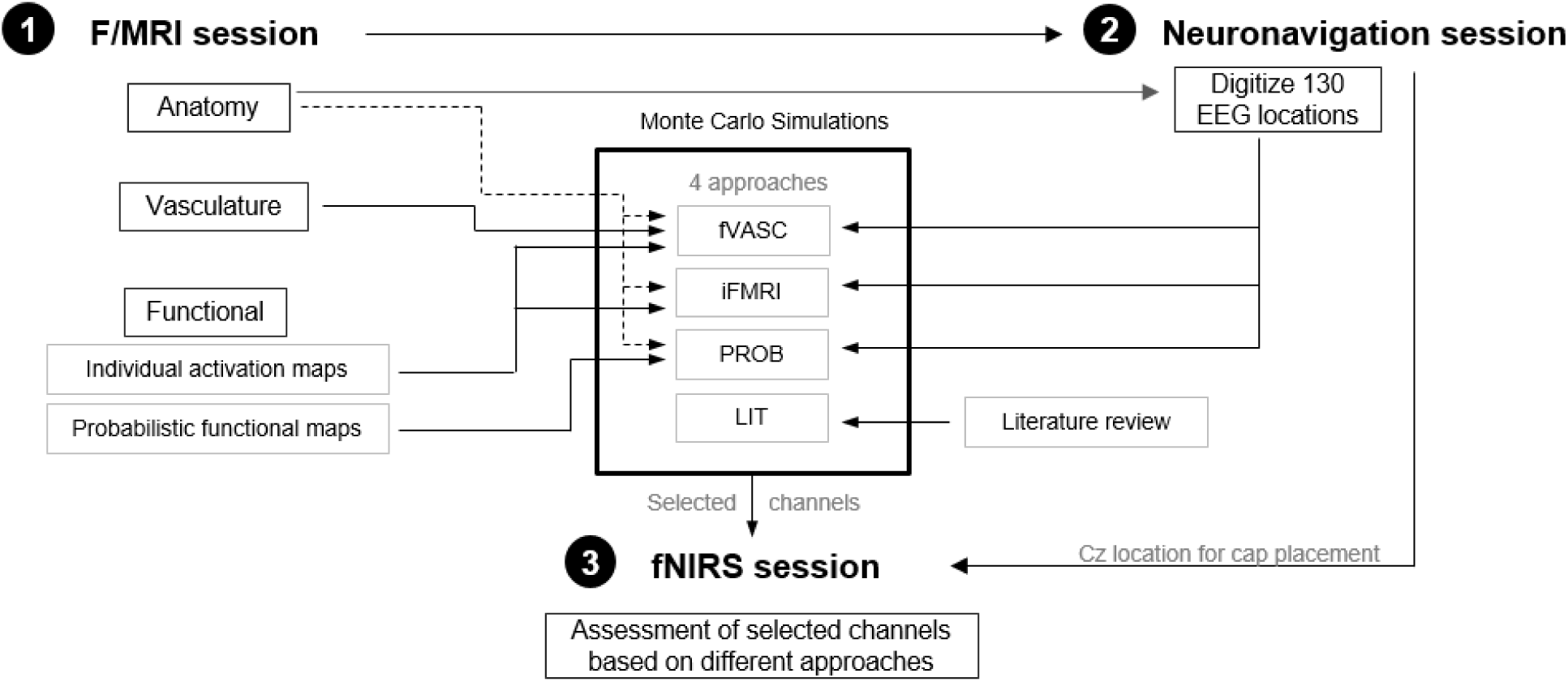
Overview of the present study. The study consisted of three separate sessions: one (f)MRI, one neuronavigation and one fNIRS session. The first two sessions aimed at collecting necessary information to create the different optode layouts for each participant. Specifically, the LIT approach used a literature review to design the optode layout. The PROB approach used probabilistic functional MRI maps, individual anatomical data and head-anatomy information for channel selection. The iFMRI approach used individual anatomical data and individual functional activation maps, together with head-anatomy information for channel selection. Finally, the fVASC approach used individual anatomical, functional and vascular data, together with head-anatomy information for channel selection. Monte Carlo simulations were used to select the best channel pair for each approach, mental-imagery task and participant. The selected channels were used during the fNIRS session to obtain information on signal quality and to measure functional activity elicited by the mental-imagery tasks.

Twenty-one participants (eleven females) were recruited for the f/MRI session. From these participants, seventeen (eleven females) took part in the neuronavigation session and sixteen (ten females) participated in the fNIRS session (see Table 1 for a summary) as some participants became unavailable over the sessions. Participants did not have a history of neurological disease and had a normal or corrected-to-normal vision. The experiment conformed to the *Declaration of Helsinki* and was approved by the ethics committee of the *Faculty of Psychology and Neuroscience*, *Maastricht University*. Informed consent was obtained from each participant before starting the experiment. Participants received financial compensation after each session.

**Table 1.** Summary of participants’ characteristics and involvement of different experimental sessions.

| Participant ID | f/MRI<br>(N=21) | Neuronavigation<br>(N=17) | fNIRS<br>(N=16) | Gender | Age range | Handedness |
| --- | --- | --- | --- | --- | --- | --- |
| **P01** | YES | YES | YES | Female | 25-30 | Left |
| **P02** | YES | YES | YES | Female | 25-30 | Right |
| **P03** | YES | YES | YES | Male | 25-30 | Left |
| **P04** | YES | YES | YES | Female | 25-30 | Right |
| **P05** | YES | YES | YES | Female | 25-30 | Right |
| **P06** | YES | YES | YES | Female | 40-45 | Right |
| **P07** | YES | **NO** | **NO** | Male | 25-30 | Right |
| **P08** | YES | **NO** | **NO** | Male | 30-35 | Right |
| **P09** | YES | YES | YES | Male | 25-30 | Right |
| **P10** | YES | YES | YES | Female | 25-30 | Left |
| **P11** | YES | YES | YES | Female | 25-30 | Right |
| **P12** | YES | YES | **NO** | Female | 25-30 | Right |
| **P13** | YES | **NO** | **NO** | Male | 20-25 | Right |
| **P14** | YES | YES | YES | Male | 30-35 | Right |
| **P15** | YES | YES | YES | Male | 30-35 | Right |
| **P16** | YES | YES | YES | Male | 25-30 | Right |
| **P17** | YES | YES | YES | Female | 20-25 | Right |
| **P18** | YES | **NO** | **NO** | Male | 25-30 | Right |
| **P19** | YES | YES | YES | Female | 20-25 | Left |
| **P20** | YES | YES | YES | Female | 20-25 | Right |
| **P21** | YES | YES | YES | Female | 25-30 | Right |

### 2.1 f/MRI session

#### 2.1.1 Data acquisition

In this one-hour long session, anatomical, functional and (brain- and scalp) vascular data were acquired at a Siemens Magnetom Prisma Fit 3 Tesla (T) scanner at the *Maastricht Brain Imaging Center*, Maastricht, The Netherlands (see Fig. 2).

**Fig 2.**
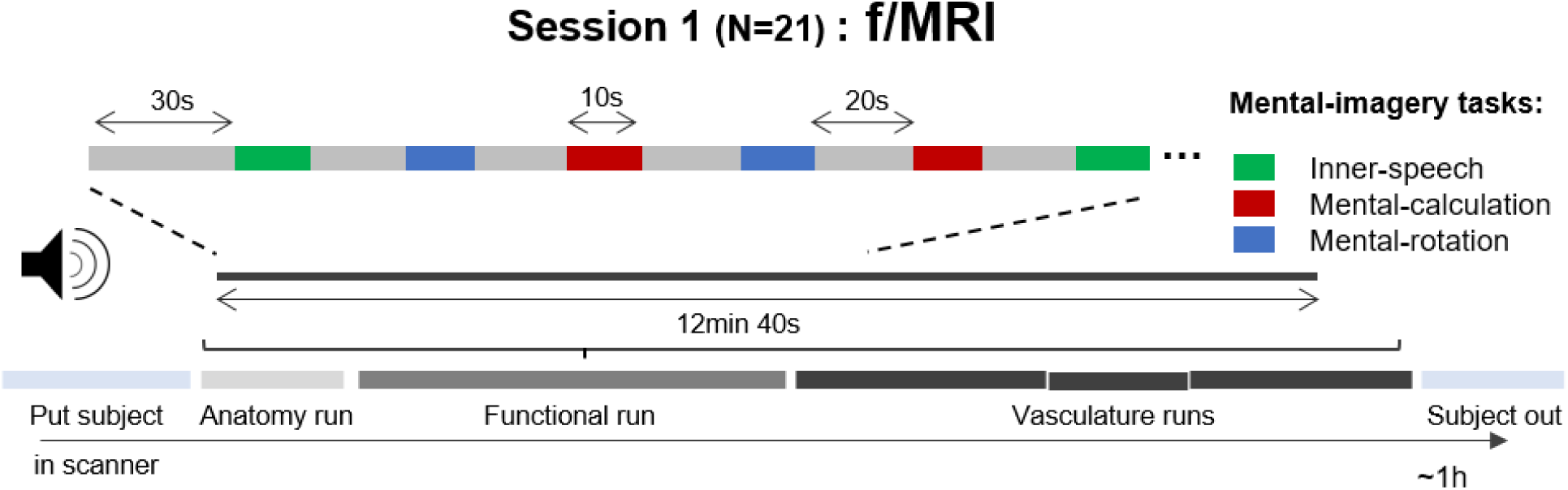
Schematic representation of session 1. Twenty-one participants underwent a one-hour long experiment in the MRI scanner, during which individual anatomical, functional and vascular data were collected. During the functional run, participants had to perform inner-speech, mental-calculation or mental-rotation for 10s each with interleaved resting periods of 20s. Task order was randomized

We used an magnetization prepared-rapid gradient echo (MPRAGE) sequence to collect structural T1-weighted MRI data, with the following parameters: repetition time (TR)=2250ms, echo time (TE)=2.21ms, inversion time (TI)=900ms, flip angle (FA)=9°, number of slices=192, 1-mm isotropic resolution, duration=5:05min. 2D Gradient Echo echo•planar imaging sequence with a TR=1s, number of slices=36, and 3-mm isotropic resolution was used to acquire functional data. Cerebral and pial vascular data was collected using 2D• and 3D• Time-of-Flight (TOF) sequences (FA=60°/18°, TR=21ms/20ms, TE=4.83/3.3ms, number of slabs=1/5, number of slices in slab=75/40, with distance factor=-33/-20%, 0.7-mm isotropic resolution, duration=9:11/4:56min). Finally, scalp-vascular data was obtained with a Multi•Echo Gradient Echo (GE) sequence with four different echoes (TR=34ms, TE_1_/TE_2_/TE_3_/TE_4_=3.02/8.56/15.11/23.91ms, number of slices=192, 0.7-mm isotropic resolution, duration= 8:06min).

#### 2.1.2 Experimental design

Participants performed one ∼13-min long functional run, where they were acoustically cued to rest (“Rest”) or perform one of the three mental-imagery tasks, namely inner- (covert) speech (“Speech”), mental-calculation (“Calculate”) or mental-rotation (“Rotate”). The order of the task trials (eight per mental task) was randomized. They were instructed to covertly recite a text they knew by heart (*e.g.*, a poem) when they heard “Speech”. Participants were asked to calculate multiplication tables of multiples of 7, 8, or 9 up to the decuple when they heard “Calculate”. When they heard “Rotate”, participants had to imagine a diver jumping from a tower into the water while he spins around several times in the air. Participants were trained on the tasks for approximately 10min before entering the MRI scanner. During training, they had to recite overtly the chosen text and the multiplication tables for the inner-speech and mental-calculation tasks, respectively to ensure the speed was consistent, and to repeat the same procedure covertly until they felt comfortable with the tasks. As for the mental-rotation task, participants watched short clips of a jumping diver until they could comfortably imagine the movement. We instructed participants to perform the mental-imagery tasks, which lasted 10s, until they heard the instruction “Rest”. During resting period, participants were requested not to do any specific mental activity and not to do/think about anything in particular for 20s (see Fig. 2 for a visualization of the run). Participants were asked to keep their eyes closed throughout the functional run. After the session, participants’ strategies were noted down and saved for the fNIRS session. BrainStim v1.1.0.1 stimuli presentation software (Gijsen, S., Maastricht University, The Netherlands) was used for both, the f/MRI and fNIRS sessions.

#### 2.1.3 Data analysis

Unless stated otherwise, all f/MRI data analyses were performed in BrainVoyager QX v2.8 (Brain Innovation B.V., Maastricht, Netherlands).

##### 2.1.3.1 Structural data

Structural images were aligned to the plane containing the anterior and posterior commissures, corrected for spatial-intensity inhomogeneities and brain-masked. The white/grey matter (WM/GM) and grey matter/cerebrospinal (GM/CSF) boundaries were detected using automatic segmentation tools. These images were inspected, manually corrected when necessary and used to create WM and GM reconstructions of the cortical surface. In addition, the (head) skin surface was automatically segmented and reconstructed. These reconstructions were used for the neuronavigation session (see below).

Cortex-based alignment (CBA) is a whole-cortex alignment scheme ^34–37^ which uses curvature information of the cortical surface to iteratively reduce misalignment across participants and in turn increase functional overlap on the group level ^38^. We used this approach to define our probabilistic functional maps. For that, individual WM reconstructions of each hemisphere were aligned to a dynamically generated group average (N=21).

##### 2.1.3.2 Functional data

Data were pre-processed using inter-scan slice-time correction, 3D rigid-body motion correction (applying Trilinear interpolation for detection/sinc interpolation, for correction), and temporal high-pass filtering with a general linear model (GLM) Fourier basis set of 3 cycles/run. Functional data of 3-mm iso-voxel resolution were spatially co-registered to the structural image by using a gradient-based intensity-driven fine-tuning alignment.

###### Generation of individual functional maps

We first calculated a voxel-wise GLM. The model contained a separate boxcar predictor for each of the mental-imagery task conditions convolved with a standard double-gamma hemodynamic response function (onset time=0s, response undershoot ratio/time to response peak=6s/6s, time to undershoot peak=16s, response/undershoot dispersion=1s/1s), and six additional predictors estimated from the motion-estimation procedure in BrainVoyager QX (translation and rotation in x, y and z direction). Individual functional maps were created in volume space by contrasting the particular mental-imagery task predictor *vs.* the rest condition (for each of the three tasks separately) in the voxels that were part of the fNIRS-coverage mask. This mask was created to mask out active voxels from deeper regions, as we did not expect the fNIRS signal to be sensitive to these regions ^39^, see supplementary materials Sec. A.1 and Fig. S1 for details. Activation maps were corrected using a cluster threshold that allowed for a 5-% loss of active voxels. These functional maps were then sampled to surface activation maps (from −1mm to +3mm from the GM/WM segmentation boundary).

###### Generation of probabilistic maps

While it is not uncommon for researchers to have previously acquired anatomical MRI data of the same participant ^38, 40^, having individual anatomical *and* functional data of the same participant represents a less likely scenario ^38^. In the absence of individual functional data, probabilistic functional maps can be generated from other individuals whose functional data are available.

Probabilistic functional maps were created separately for each participant and mental-imagery task following a leave-one-subject-out procedure ^41^. For each participant, surface activation maps from the remaining participants were aligned using individual transformation files derived from the CBA approach. It should be noted that MR *vs.* Rest map from P08 was excluded from subsequent analyses as the participant reported not being able to perform the mental-imagery task correctly and having used an alternative cognitive strategy instead. Thus, the probabilistic maps for each participant were created based on N=20 participants for the IS and MC tasks and based on N=19 participants for the MR task. We discarded mesh vertices that were active in less than 20% of the sample size for each task and hemisphere. The resulting probabilistic maps for each hemisphere were transformed back into individual volume space (by interpolating from −1mm to +3mm from the GM/WM segmentation boundary) and smoothed with a 2mm full-width-half-maximum kernel. The final maps (three per participant) were used as region of interests for Monte Carlo simulations (see Sec. 2.3.2). Examples of probabilistic maps are shown in Fig S2.

##### 2.1.3.1 Vascular data

###### Cerebral and Pial vasculature

2D and 3D TOF data were aligned individually to an up-sampled version (0.7-mm isotropic resolution) of the anatomical data of the same session for each participant, following the same co-registration approach as for functional maps described above. Vascular data were segmented with automatic segmentation tools in BrainVoyager QX (intensity-based segmentation) and the software *Segmentator* (intensity gradient-based segmentation ^42^) and manually corrected when necessary. The latter was done using ITK-snap ^43^ and BrainVoyager QX. The segmented vascular structures from 2D and 3D TOF data were then combined and were down-sampled to 1-mm isotropic resolution. The analyses procedures are summarized in a flow-chart diagram (Fig. S3) and an example reconstruction is shown in Fig. S4.

###### Scalp vasculature

All four echo images derived from the multi-echo GE protocol were first aligned to the 0.7-mm isotropic resolution anatomical images for each participant. We then isolated the extracerebral tissues by masking out the brain using *FSL BET* v5.0 ^44^. Depending on which image(s) showed higher contrast for vascular structures, segmentation was performed manually in BrainVoyager QX using a combination of the four echoes or using the later echo images, *i.e.*, TE_3_=15ms and TE_4_=23ms, which showed higher contrast for vascular structures than earlier echoes. The segmented vascular structures were then down-sampled to 1-mm isotropic resolution. The analyses procedures are also summarized in the flow-chart diagram provided in Fig. S3 and an example reconstruction is shown in Fig. S4.

### 2.2 Neuronavigation session

Seventeen of the originally included 21 participants underwent this session, as P07, P08, P13 and P18 dropped out of the study. A neuronavigation system (Zebris CMS20 ultrasound system, Zebris Medical GmbH, Isny, Germany) in combination with BrainVoyager QX 2.1 TMS Neuronavigator software (Brain Innovation, Maastricht, Netherlands) was used to acquire the coordinates of 130 EEG positions for each participant (see Fig. 3). These 130 locations were determined based on the layout of EasyCap 128Ch ActiCap (EasyCap GmbH, Herrsching, Germany) whose size was selected based on individual head sizes. First, the head circumference for each participant was measured using a measuring tape. The cap was placed on and was secured using a chin band. Next, its position was adjusted so that the Cz location would be exactly half the nasion-inion distance. The inion was defined as the top part of the pronounced structure in the occipital region. In order to ensure that the cap was not tilted or shifted to one side, the distance between the left and right pre-auricular points was measured and the cap was gently moved in this virtual coronal plane until Cz was located half this distance. The preauricular points were defined as the location where the mandibular bone moves with the opening and closing of the mouth. Finally, the cap was secured with medical tape on the forehead to prevent any unwanted cap shift. The Cz location details (in terms of nasion-inion and pre-auricular distance) together with the cap size were noted down for the fNIRS session.

**Fig 3.**
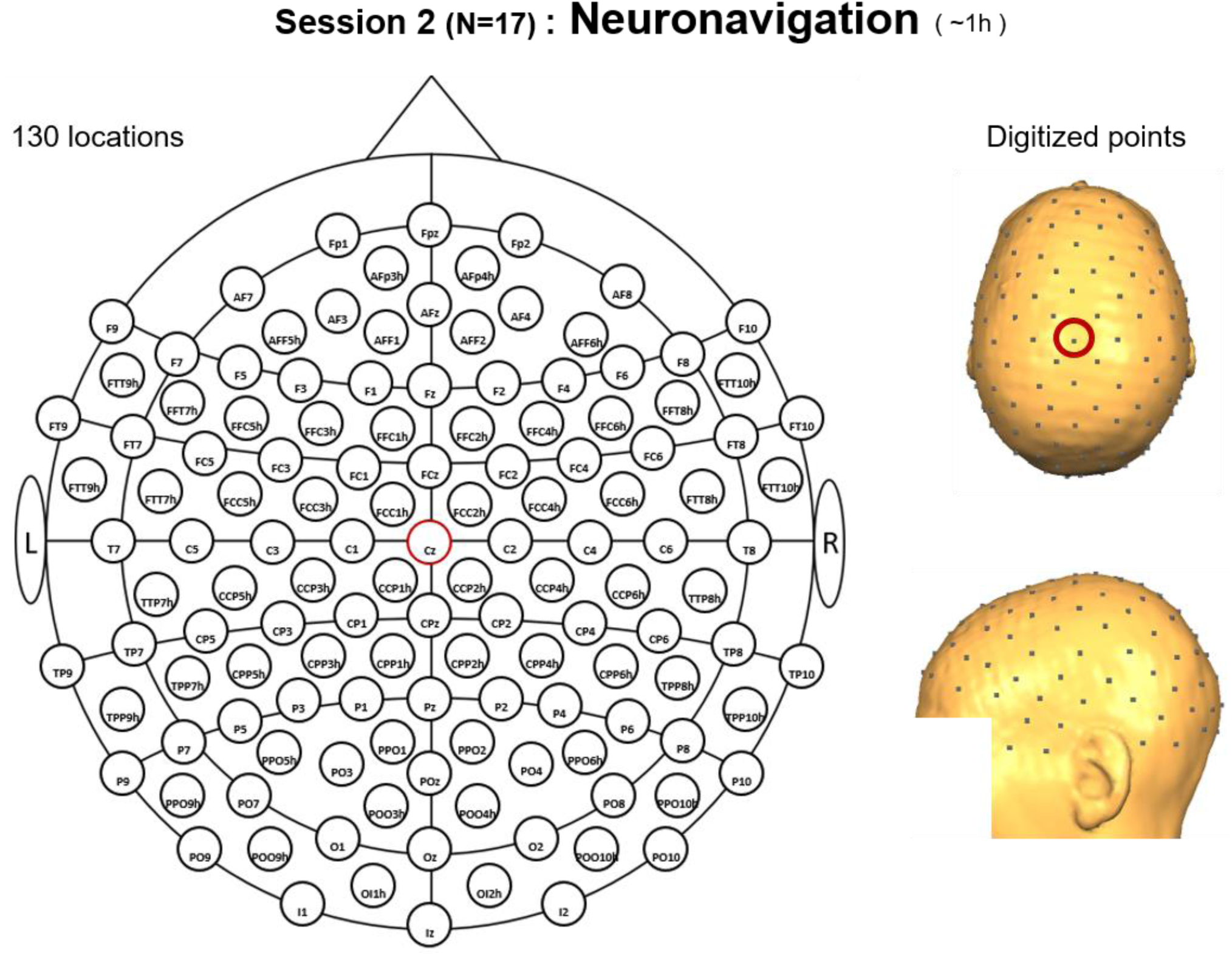
Schematic (left) and reconstructed (right) locations recorded during the Neuronavigation session. This layout is an extension of the international 10-20 system, it contains 130 locations and the nomenclature is based on ^20^. The Cz location is indicated with a red circle. The schematic representation is based on the NIRx montage editor template, while the reconstructed locations belong to participant P04.

Single ultrasound markers (three in total) were attached to the participant’s head using adhesive stickers. Next, three reference points (inion and left and right preauricular points) defined on the participant’s head were used for the co-registration of the structural MRI image with the participant’s head in the external (real) world. Once these steps were completed, the 130 EEG locations marked on the cap were digitized. The session lasted approximately 1h.

### 2.3 fNIRS session

#### 2.3.1 Participants

P12 dropped out of the study. Thus, 16 of the 17 participants that participated in the fMRI and neuronavigation sessions took part in this session, out of which ten were female (mean age=29.81±5.22).

#### 2.3.2 Designing approach-specific optode layouts

This process can be divided into three main stages: channel sensitivity computation, channel selection and building a participant-specific layout (see Fig. 4 for a summary). The first stage aimed at computing the channel-sensitivity profiles using Monte Carlo simulations. Each of the four approaches had a unique combination of ROI definition/type, software and brain model used to compute the simulations. During the second stage, the most-informative channels were selected for each of the four approaches, based on the solution to an optimization problem subject to a set of constraints. The first and second stages were repeated until approach- and task-specific optode layouts were created (twelve per participant, since there were three tasks and four approaches). The last stage aimed at combining all optode layouts into a single one individually for each participant.

**Fig 4.**
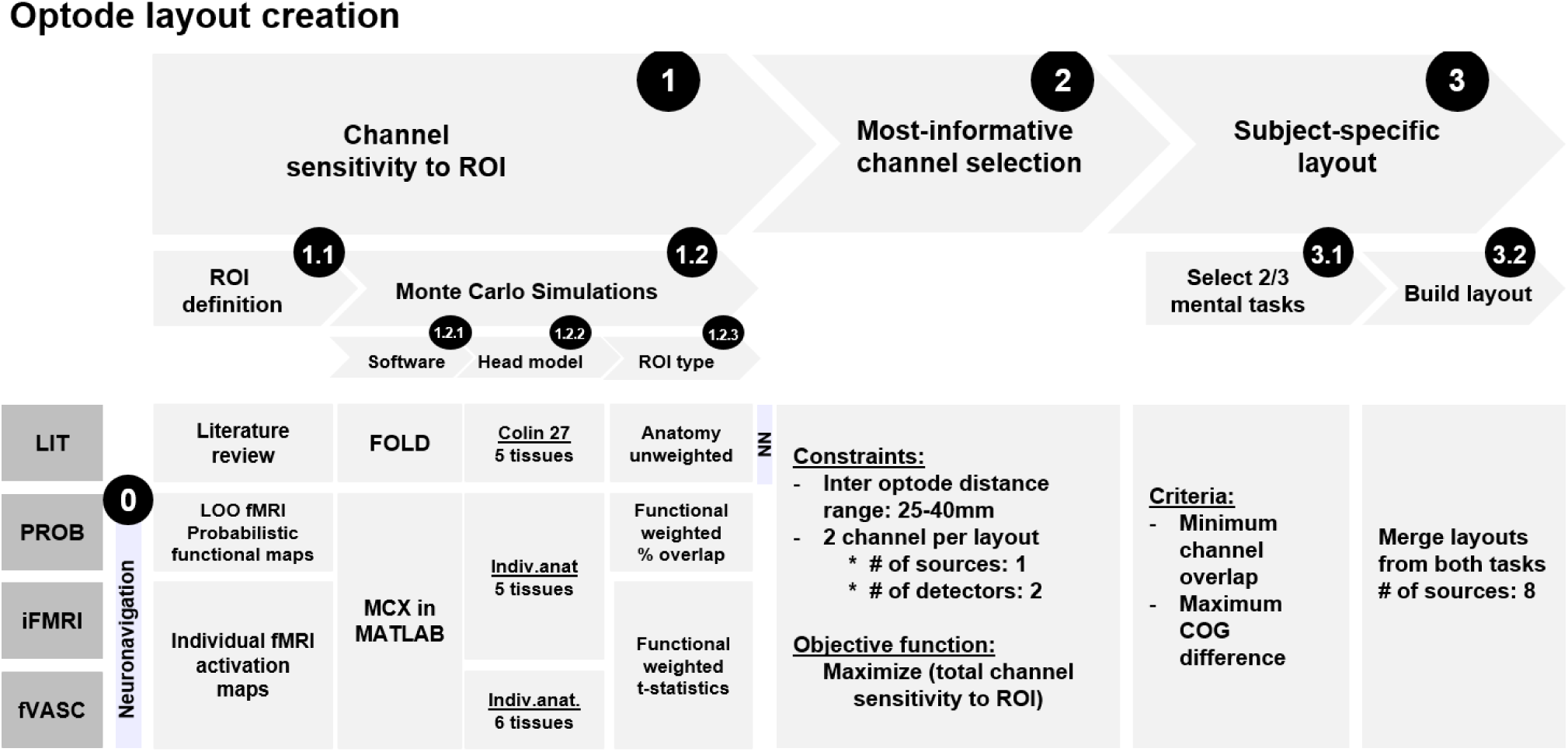
Summary of the key steps involved in optode-layout design for each of the four approaches evaluated in the present study. The process was divided into three main stages: (1) channel sensitivity to ROI computation, (2) channel selection and (3) building a subject-specific layout. For the first stage, each of the four approaches had a unique combination of ROI definition/type, software and brain model used to compute the Monte Carlo simulations. During the second stage, the most-informative channels were selected for each of the four approaches and two mental-imagery tasks. The last stage combined all the layouts into one. LOO = leave-one-out; COG = center of gravity; NN = neuronavigation.

##### 2.3.2.1 Channel sensitivity to ROI computation

All four approaches (LIT, PROB, iFMRI, fVASC) were based on the light sensitivity profiles to a given ROI, but they differed in the following aspects (see Table 2 for a summary):

1. <u>Software for Monte Carlo simulations</u> The LIT approach represents a scenario where no individual MRI anatomical data is available and the target ROI is selected based on a literature review. Given such scenario, FOLD toolbox ^30^ provides an easy way to compute the sensitivity profiles to the selected ROIs. This is because FOLD uses atlas head models as inputs to the Monte Carlo simulation and offers different brain parcellation atlases for ROI definition in the target head-model space. In addition, it is freely available, easy to install and has a user-friendly graphical interface. FOLD uses MCX package ^45^ to compute the light sensitivity profiles of optodes placed in pre-defined locations on the scalp, namely points corresponding to the extended 10-10 and 10-5 systems (130 points in total). It then provides a list of channels with the highest sensitivity to the ROI that can be exported for subsequent computations. PROB, iFMRI and fVASC approaches represent scenarios where individual MRI anatomical data are accessible. Since FOLD does not offer the option of using individual head models to compute Monte Carlo simulations, these were computed using the MCX package directly through its MATLAB interface (v2015b, The MathWorks, Inc., Natick, Massachusetts, United States).
2. <u>Head models and tissue segmentations</u> Monte Carlo simulations require the anatomical head models to be segmented into different tissues. This is necessary for photon-transport simulations as different tissues of the human head present different optical properties (absorption, scattering, anisotropy and refraction). For the LIT approach, we used the MNI Colin27 head atlas (the default atlas available in FOLD). FOLD uses a five-layer segmentation of the MNI Colin27, which consists of scalp, skull, CSF, GM and WM tissues. For the remaining approaches, a five-layered model was created from the individual anatomical images using a hybrid segmentation algorithm ^40^. This algorithm, developed in MATLAB and available upon request from the authors, takes as input the standard GM and WM segmentations of a T1-weighted image from FreeSurfer and applies sequential morphological operations implemented in iso2mesh tools to accurately reconstruct skull, scalp, and CSF layer thickness. The GM and WM segmentation images were created in FreeSurfer v06 ^46^ using the standard processing stream (*recon –all*, which took ∼10 h per participant). The resulting tissues from the hybrid segmentation algorithm were converted into compatible BrainVoyager QX files to visually inspect and manually correct them if necessary. Although GM and WM segmentation files had been created in BrainVoyager QX in a previous step (see Sec. 2.1.3.1), the automatic segmentation in BrainVoyager usually disregards the cerebellum. We thus used the segmentations from FreeSurfer to create a head model for Monte Carlo simulations. From the corrected segmentation files, a single image file was created by assigning integer values ranging from 1 to 5 to the different tissues (as in FOLD). Specifically, voxels corresponding to scalp were assigned the value 1, voxels corresponding to skull were assigned value 2, CSF 3, GM 4 and WM 5. The remaining voxels were assigned value 0 (air). We ensured that voxels inside the head were not assigned the value 0 by first identifying them and subsequently assigning the value dictated by their direct neighbors. The fVASC approach differed from the PROB-based and the iFMRI-based approaches in that vascular structures were included in the head model. For that, both pial/brain and scalp vasculature segmentations were combined and included as the sixth layer. To prevent voxels being assigned to two different tissues simultaneously, all voxels considered as vascular tissue were removed from the remaining five tissues. Importantly, our segmentations could not distinguish veins from arteries and all voxels were treated as veins. Both, five- and six-layered models are shown in Fig. S5.
3. <u>Optical properties</u> For comparability purposes across approaches, we used the average optical properties across four NIRS wavelengths (690, 750, 780 and 830nm) as in FOLD. We defined the optical properties of vascular structures based on the scattering, absorption and anisotropy values provided by ^31^. We refer the reader to the Supplementary Tables S1 and S2 for computation details and Table 2 summary table of the optical properties used in the present study.
4. <u>ROI selection and definition</u> The ROIs for the LIT approach were selected based on a literature review of the three mental-imagery tasks used in this study (we refer the reader to the supplementary materials Sec. A.2 and Tables S3 and S4 for a summary of the reviewed studies and the selected ROIs, respectively). These ROIs were defined in the MNI Colin27 brain based on the Jülich histological atlas available in FOLD. The selected ROIs for the PROB-based approach were the active regions of the individual probabilistic mental-imagery maps. For iFMRI and the fVASC approaches, individual mental-imagery contrast maps were used as ROIs (see Sec. 2.1.3.2).
5. <u>Inter-optode distance</u> FOLD performs the Monte Carlo simulations on neighboring optical positions of 10-10/10-5 systems only (that have a median inter-optode distance of 36mm) to avoid too long distances that cannot provide measurements with a proper signal-to-noise ratio ^30^. For PROB, iFMRI and fVASC approaches, we only considered channels whose inter-optode distance was in the range of 20-45mm for Monte Carlo simulations. The number of channels differed across participants as the inter-optode distance could differ with varying head size/shapes across participants (see Table 3 for participant’s cap size).
6. <u>Computation of the sensitivity of a channel to a given ROI</u> Monte Carlo simulations are used to calculate the fluence distribution produced by a source transmitting light into a highly scattering medium ^39^. By taking the product of the source and detector fluence distributions (also known as adjoint field), the photon measurement density function can be calculated ^47^. This is equivalent to the light sensitivity profiles mentioned earlier. FOLD calculates channel-wise normalized sensitivity profiles from the adjoint field by scaling the adjoint field with the sum of sensitivity of all voxels, so that each voxel represents percentage sensitivity to the whole volume. Then, the sensitivity of a channel to a given ROI is computed as a weighted mean of the voxels within the ROI to the sensitivity of voxels corresponding to the brain (GM and WM):

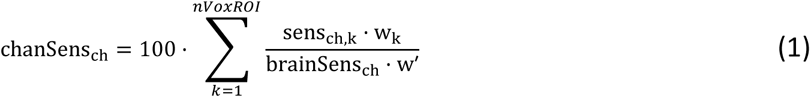

where *nVoxROI* corresponds to the number of voxels comprising the target ROI, sens_ch,k_ is the normalized sensitivity value for channel *ch* and voxel *k*, brainSens_ch_ is the normalized sensitivity of channel *ch* of all GM and WM voxels, and *w* corresponds to the value (weight) of the voxel *k* in the target ROI (adapted from ^30^). The four approaches differed in the *nVoxROI* and the *w* parameters. The LIT approach assumed that all voxels belonging to a particular (anatomical) ROI contributed equally to the computation of the sensitivity of a channel to a given ROI and thus all weights were set to one. The PROB approach used probabilistic functional maps that represent the percent overlap of voxels across participants and thus weights ranged between 0 and 100%. As for iFMRI and fVASC, they relied on individual functional activation maps whose weights represent *t*-statistic values and ranged between 0 and 15.

**Table 2.**
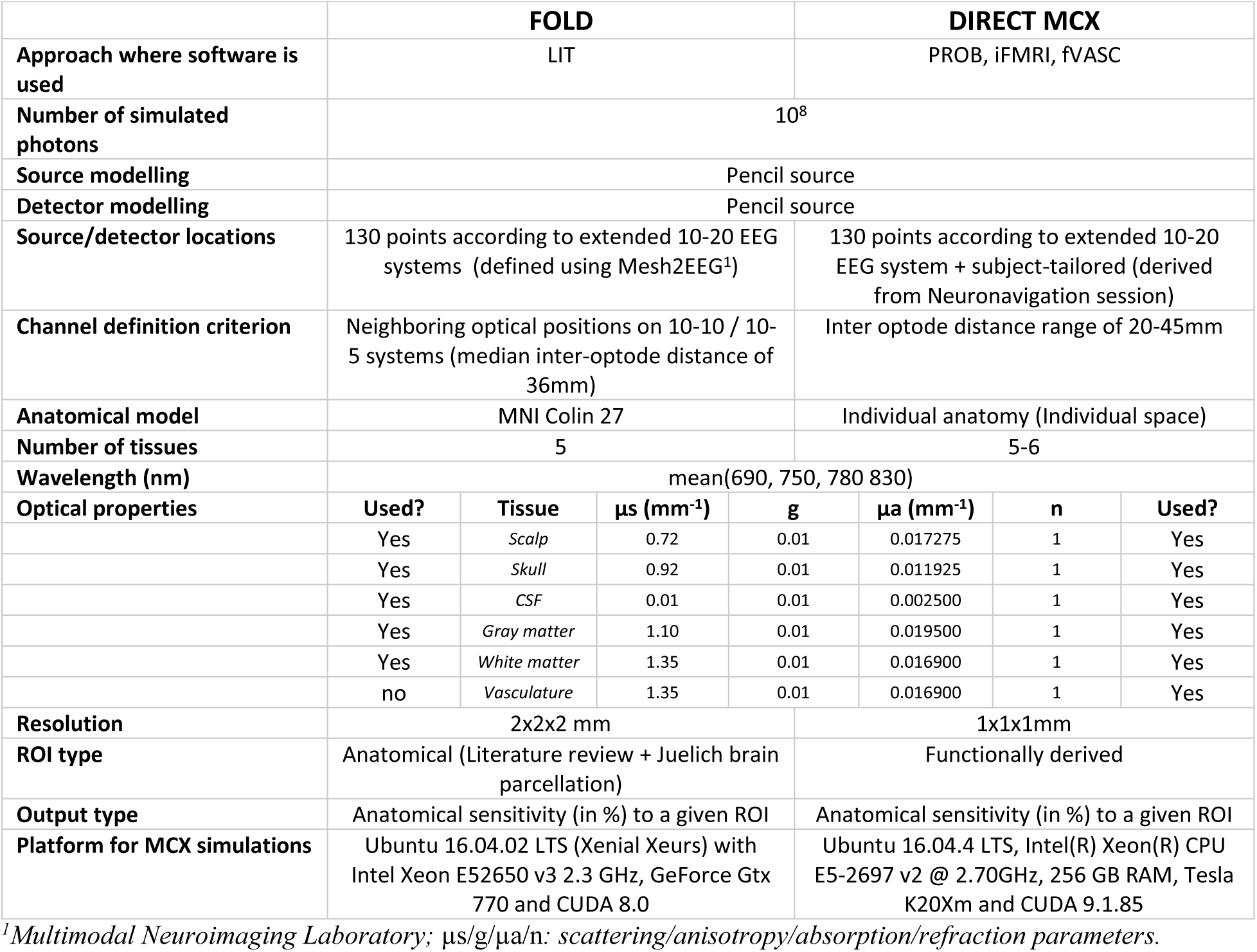
Comparison between Monte Carlo simulation approaches.

**Table 3.** Subject-specific fNIRS-session summary and optode-layout information.

| Participant ID | Cap Size (cm) | Mental tasks |  | # Runs | # Optodes S \| D |  | # NDC channels | IOD (mm) |  |
| --- | --- | --- | --- | --- | --- | --- | --- | --- | --- |
|  |  |  |  |  |  |  |  | Mean | Std. dev. |
| P01 | 56 | MC | MR | 6 | 8 | 7 | 13 | 29,92 | 2,60 |
| P02 | 56 | IS | MC | 6 | 8 | 9 | 16 | 29,00 | 3,48 |
| P03* | 58 | IS | MR | 6 | 8 | 6 | 16 | 34,06 | 6,84 |
| P04 | 56 | MC | MR | 6 | 8 | 10 | 15 | 31,60 | 4,79 |
| P05 | 60 | IS | MC | 6 | 8 | 4 | 12 | 30,67 | 4,52 |
| P06 | 56 | MC | MR | 6 | 8 | 9 | 14 | 31,07 | 4,20 |
| P09 | 60 | MC | MR | 6 | 8 | 5 | 9 | 29,33 | 4,42 |
| P10 | 56 | MC | MR | 6 | 8 | 7 | 10 | 30,40 | 4,12 |
| P11 | 56 | MC | MR | 6 | 8 | 4 | 12 | 30,17 | 3,19 |
| P14 | 58 | MC | MR | 5 | 8 | 6 | 11 | 31,09 | 3,14 |
| P15 | 58 | MC | MR | 5 | 8 | 5 | 10 | 31,70 | 4,64 |
| P16 | 58 | MC | MR | 6 | 8 | 8 | 12 | 31,58 | 3,42 |
| P17 | 56 | MC | MR | 6 | 8 | 5 | 11 | 30,18 | 4,33 |
| P19 | 56 | MC | MR | 6 | 8 | 7 | 12 | 29,83 | 2,79 |
| P20 | 56 | MC | MR | 6 | 8 | 10 | 15 | 31,13 | 4,50 |
| P21* | 54 | IS | MR | 6 | 8 | 10 | 14 | 29,07 | 3,25 |

Note: P03 and P21 were excluded from data analysis (see participant exclusion criteria)
Abbreviations: NDC= normal distance channels; IOD= inter-optode distance; MC = mental-calculation; MR= mental- rotation.

For the LIT approach, channel sensitivity to a given ROI was computed separately for 10-10 and 10-5 systems as they cannot be computed simultaneously in FOLD. FOLD allows choosing the minimum value of the channel sensitivity to a given ROI to select/discard channels. We set this threshold to 0% in order to select all channels that were somewhat sensitive to the target ROI and combined the list of output channels for every ROIs that was used for each mental-imagery task. If a channel appeared multiple times for a task, we selected the highest sensitivity value among all instances. As for the remaining three approaches, all channels that were considered for the Monte Carlo simulations together with their associated sensitivity values were selected as input to the next step.

##### 2.3.2.2 Optimization of the optode layout

We determined the most informative set of channels (separately for each approach and mental-imagery tasks) by maximizing their total sensitivity to the target ROI. The maximization problem was subject to two constraints:

1) The inter-optode distance did was limited to the 25-40mm range. We used individual inter-optode distance measures derived from the neuronavigation session for this step. It is important to note that this was applied to all four layouts (thus including the layout based on the LIT approach). The FOLD toolbox (used for LIT approach) uses near-neighbor channels with a median inter-optode distance of all channels to be 36mm, in MNI space^30^. We used this additional information to ensure that (1) all channels were in the 25-40mm range in the subject-specific space, and that (2) the signal-quality standards for all approaches were as similar as possible.

2) The optode layout for each approach consisted of two channels that shared a common detector (thus including three optodes per approach). Since we did not distinguish between sources and detectors in the Monte Carlo simulations, it is important to realize that the sensitivity of the channel will remain the same whether one considers optode X a source and optode Y a detector, or vice-versa. However, due to the second constraint, the algorithm may select a different channel pair that maximizes the total sensitivity to the ROI depending on which optode is considered a source or a detector. To ensure that as many candidate channels as possible were considered during the optimization approach, the optimization problem was solved twice: (1) using the original channel pool that consisted of all optode pairs that were considered for the Monte Carlo simulations (on average, there were 633.25 channels [SD=44.13] across participants); (2) considering their swapped versions (sources were considered detectors and vice-versa).

We followed an iterative approach to address the optimization problem. It begins with the construction of an empty solution, where no optode pair is selected. The algorithm then prunes the optode pairs that do not satisfy the inter-optode distance range constraint. Next, the algorithm ranks all possible optode pairs according to their contribution to the total sensitivity and selects one pair as the seed in each iteration. The algorithm then transfers the selected optode pair to the solution matrix and it removes from the list the channels that do not share the same detector. Next, it selects the first channel from this list (*i.e.,* the one with the highest sensitivity). Since the target number of channels (=2) has been reached after this step, the accumulated total sensitivity of the selected two channels and the source-detector indices are stored in the solution matrix. These steps are repeated until all optode pairs are used as seeds. Finally, the two channels that lead to the highest total sensitivity for either constraint set constitute the selected channels for creating the setup.

##### 2.3.2.3 Creating the setup

###### Mental-imagery task selection

Two out of the three mental-imagery tasks that participants performed during the f/MRI session were selected for the fNIRS session. This measure was necessary as pilot measurements performed with optode layouts designed to account for all three tasks elicited high discomfort in participants. This decision ensured that the optode setup would maximally consist of 24 optodes (3 optodes per layout × 4 approaches × 2 motor-imagery tasks), which should constitute a reasonably comfortable setup for participants and thus should prevent them from withdrawing from fNIRS recordings due to setup-related discomfort ^48–50^ This selection was carried out at the individual subject level. For that, we first calculated the number of overlapping channels across all four layouts for each mental-imagery task, and selected the two tasks with the least number of overlapping channels. An additional step was used in case this approach was not sufficient to select the two tasks, where we computed the center of gravity (COG) for all four layouts per mental-imagery task and calculated the distance between COGs. The tasks with the least number of overlapping channels and highest distance between them were the selected tasks. See Supplementary Table S5 for a summary of the mental-imagery task selection procedure and Table 3 for the resulting selected task pair per participant.

###### Combining all channels into a single layout

The eight layouts (four per task) were combined manually into a single one. This was first carried out digitally to simulate the final arrangement using schematic representations of source and detector positions. It consisted of two steps: an initial step combined all four layouts for each mental-imagery tasks and both layouts were combined into one in the second step. It could be that the source-detector arrangement was not compatible across layouts (within or across mental tasks), since a source in a given channel cannot be a detector in another one (or *vice versa*). To account for such possibility, we first swapped sources for detectors in the problematic spots. This step solved the compatibility problem in all but four participants (P05, P16, P17 and P19). For these participants, using a different mental-imagery task combination solved the issue (see Supplementary Table S5). Since the fNIRS system used in this study uses lighter wires for sources than for detectors, we rearranged sources and detector positions in all participants (when possible) to maximize the number of sources while preserving the channels defined in the optimization step. It is important to note that each participant ended up with a unique optode layout, with a varying number of optodes (see Table 3 and Fig. S6).

#### 2.3.3 Experimental design

The fNIRS experiment consisted of one session that lasted approximately 1.5h. During this time, participants performed six, around 8-min long functional runs. In each of the runs, participants were acoustically cued to perform one of the two mental-imagery tasks selected for them or to rest. Six, 10-s long trials were presented for each mental-imagery task, interleaved with a jittered rest condition with mean duration of 22s (jittering was of ± 2s), see Fig. 5. Thus, participants performed 60 trials for each mental-imagery task across the six runs. Trials were pseudo-randomized across runs. Participants were instructed to use the same strategy they used in the scanner (first session). For that, participants were given a document prior to the fNIRS experiment where their strategies had been noted down. Participants were asked to avoid any potential jaw movements during the functional runs and to keep their eyes closed throughout the run.

**Fig 5.**
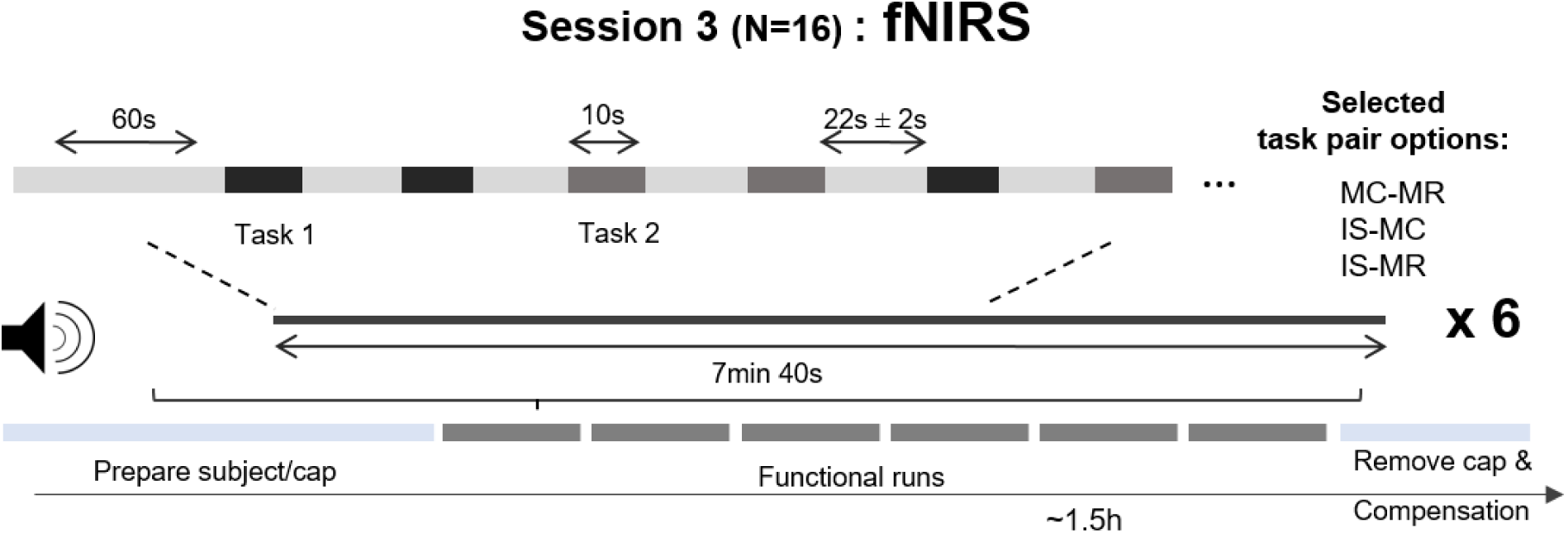
Schematic representation of a functional run during the fNIRS session. During each mental-task period, participants were acoustically cued to perform one of the two mental-imagery tasks for 10s while keeping their eyes closed. When participants heard “rest”, they were asked to stop the task and await the next instruction. Abbreviations: IS= inner-speech; MC = mental-calculation; MR= mental-rotation.

#### 2.3.4 fNIRS signal acquisition

fNIRS data were recorded using a continuous-wave system (NIRScout-816, NIRx, Medizintechnik GmbH, Berlin, Germany). The optode setup varied across participants, but they had some features in common: all setups contained eight sources and eight short-distance channels (SDC). The SDCs were formed by short-distance detectors placed at 8mm from a given source. The inter-optode distance of the standard channels (here on called normal-distance channels, NDC) ranged from 25-40mm. Sources emitted light at wavelengths 760nm and 850nm, and the light intensity acquired at the detector side was sampled at 7.8125Hz. The fNIRS cap was placed for each participant according to the measurements taken during the neuronavigation session. Besides the standard cap fixation (using the chin band), the fNIRS cap (EasyCap 128Ch ActiCap, EasyCap GmbH, Herrsching, Germany) was fixated onto the participant’s head with three medical tape stripes (connecting the cap and the participant’s forehead) to assure the cap would not shift during the measurements. In addition, a black, plastic overcap was placed on top of the fNIRS cap to additionally prevent ambient light from reaching the spring-loaded optodes.

#### 2.3.5 fNIRS data analysis

##### 2.3.5.1 Participant exclusion criteria

Two of the sixteen participants, P03 and P21, were excluded from subsequent analysis for different reasons. The optode layout for P03 was created based on a different inter-optode distance range criterion than the rest of the participants (25-45mm vs. 25-40mm). This is because P03 was the first participant who participated in the fNIRS session and the original inter-optode distance range was expected to provide reasonable signal quality. However, this range proved to be suboptimal as four NDC and three SDC did not survive the coefficient of variation threshold (CV < 7.5%), a metric used to estimate the signal-to-noise ratio for each channel ^51^. Given the restricted number of channels comprising each layout, we created the layouts for the rest of the participants using a more conservative inter-optode distance range criterion (25-40mm range, see first constraint in Sec. 2.3.3) to ensure that all (or as many as possible) channels survive the CV threshold. Thus, P03 was excluded for comparability reasons. As for P21, the data was corrupted and could not be retrieved.

##### 2.3.5.2 Preprocessing

For every subject and run, the raw optical intensity data series were converted into changes in optical density (OD) values using Homer2 ^52^. CV values were calculated for the entire run for each channel and those with a CV >=7.5% were discarded from the analysis (see Fig. S7). Next, the motion detection algorithm *hmrMotionArtifactByChannel* was applied to the OD time-series to identify motion artifacts in each channel. We used the following parameters: AMPThresh=0.15, tMotion=0.5 and tMask=2. The SDThresh parameter ranged between 8 and 10 across participants. Motion artifact identification was visually assessed by experimenter AB and was manually corrected in case it was necessary. Motion artifacts were divided into spikes and baseline shifts. Baseline shifts were corrected using *hmrSplineInterp* algorithm in Homer2 (p=0.99), while *hmrMotionCorrectWavelet* algorithm in Homer2 (iqr=0.5) was used to correct for the spike artifacts only in the channels where motion artifacts had been detected (Fig. S8 summarizes the detected number of motion events per participant). Then, motion-corrected OD data were transformed to change in concentration values through the modified Beer-Lambert law with an age-specific differential path length factor for each participant ^53^.

##### 2.3.5.3 Assessment of degree of layout (dis)similarity across approaches

The first goal of this study was to assess whether the resulting optode layouts differed across approaches. To do so, for each pair of approach-specific layouts we calculated the number of overlapping channels and the Euclidian distance between their centers of gravity. These calculations were carried out for each mental-imagery task at the single-subject level and were averaged across participants afterwards. In addition, frequency maps for each approach were computed.

##### 2.3.5.4 Single-run estimates calculation

The Short Separation Regression approach (SSR ^54^) was applied on the unfiltered Δ[HbO]- and Δ[HbR]-NDC data to remove signal from extra-cerebral layers of the head. This was done for each NDC and chromophore by using the SDC closest to the NDC as the regressor. The SDC-corrected time course was used as input for the *ar_irls* algorithm in NIRS Brain AnalyzIR Toolbox ^55^. This algorithm uses an autoregressive (AR) model for correcting motion and serially correlated errors in fNIRS. The function was adapted to use the ordinary least squares method instead of the *robustfit* approach. The maximum AR model order to be considered was set to four times the sampling rate. The design matrix included the two task predictors convolved with a standard hemodynamic response function. The default hemodynamic response function from SPM12 was used (double gamma function, the onset of response and undershoot 6s and 16s, respectively, dispersion 1s, response to undershot ratio 6). The task predictor for Δ[HbR] was −1/3 of the Δ[HbO] amplitude. In addition, a set of low frequency discrete cosine terms were defined as confound predictors using the *dctmtx* function in NIRS Brain AnalyzIR Toolbox with a cut-off frequency of 0.009Hz.

##### 2.3.5.5 Multi-run ROI analysis

We combined the information from both channels comprising each layout to run an ROI analysis as described in Santosa and colleagues^55^ and expanded their procedure to account for multiple runs:

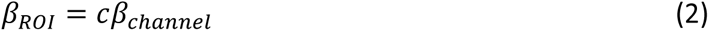

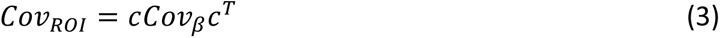

where in this study *β_channel_* is the multi-run beta estimate and the *Cov_βroi_* is the multi-run covariance matrix estimated from the concatenated residual time courses and the design matrix. Finally, *c* is the contrast vector whose coefficients are 0 if the channel does not belong to the ROI and is 0.5 in the two channels that belong to the ROI.

##### 2.3.5.6 Multi-run block averages and contrast-to-noise ratio

The SDC-corrected and unfiltered Δ[HbO] and Δ[HbR] time courses were filtered using a zero-phase, band-pass finite impulse response filter of order 1000, with cutoff frequencies of [0.008, 0.25Hz]. Block averages were computed for each channel and mental-imagery task by taking the average of all trials and runs 4s before the onset of the task until 15s after the offset of the task. The Contrast-to-Noise Ratio (CNR) as was calculated for each channel, ROI and chromophore using the formula described by ^48^:

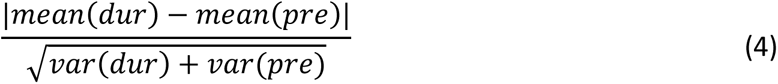

where *pre* represents the rest period from 4s before onset of task to 0s; and *dur* represents the task period from 5-15s post task-onset, as in ^56^.

##### 2.3.5.7 Statistical analysis

The second goal of this study was to compare the fNIRS-signal quality and sensitivity obtained from the optodes placed according to the four different approaches. Group differences across approaches in terms of CNR and ROI *t-*estimates were assessed using a non-parametric ANOVA (Friedman test) and follow-up Wilcoxon paired signed rank tests, one-sided and corrected for multiple comparison with the Benjamini-Hochberg method. Group differences were computed considering: (1) each mental-imagery task separately and (2) all mental-imagery tasks together. In addition, we quantified the number of participants that showed significant increase in the ROI activation.

##### 2.3.5.8 fNIRS data projection onto cortical surface and comparison with fMRI data

We used the inverse distance weighting (IDW) method described in ^57^ to interpolate fNIRS data on the cortical surface. In short, each fNIRS channel position was defined as the point in the scalp half way between the corresponding source and detector position. The cortical projection of each channel was determined by taking the point in the brain reconstruction closest to the channel position in the scalp. A sphere of radius *r* was centered in the projected cortical point and the voxels inside the sphere that were labeled as GM were assigned a weight depending on how far from the center they were located. The weight (w) was calculated as 1/d^2^, where *d* is the Euclidian distance between the projected point (center of the sphere) and a given voxel inside the sphere. At each cortical vertex *k* inside the sphere, the interpolated fNIRS data was computed as:

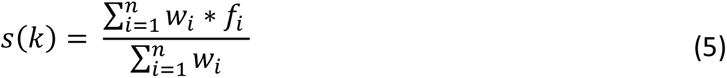

where *n* is the number of cortical projection points and *f* is the amplitude of the fNIRS channel value. Here we used two cortical projection points as two channels comprised a given layout. The channel-specific amplitude was calculated as the average value of the normalized fNIRS signal (computed as the channel time course divided by its peak value) in the range of 3s after task onset to 5s after task offset. In total, four spheres with varying radii (r = {10, 15, 20, 25} mm) were used.

We used channel-specific projection weights and projection spheres to compute spatially weighted fMRI block averages to assess the temporal correlation between fNIRS and fMRI signals. First, voxels inside the sphere of radius *r* that were labeled as GM were selected and mental imagery-specific events were extracted from each voxel’s time courses. Task-specific ROI averages were computed by weighting the contribution of each voxel according to the projection weights. The standard error of the weighted average was estimated using bootstrapping (with 100 resamples and sample size equal to 60% of the initial number of voxels). These steps were repeated for every channel across all layouts in each participant. Finally, the temporal correlations of fNIRS and fMRI block averages were computed using Spearman’s correlation.

Next to channel-specific projection weights, layout-specific projection weights were also calculated. Their computation differed in that for the latter we used the center of gravity of each layout on the scalp to determine the cortical projection point. Layout projection weights were used to extract the peak and spatially weighted mean *t*-estimates of individual fMRI activation of the voxels labeled as GM to assess how well the fNIRS ROIs targeted individual activation maps.

## 3 Results

### 3.1 Using different information sources for optode placement results in different optode-layout designs

Figure 6 shows the mean percent overlap (top panel) and mean Euclidian distance between the COGs of each pair of optode layouts across participants (bottom panel). The color of each cell indicates the standard error of the mean. The LIT approach contained no channels that overlapped with the remaining approaches for neither mental-calculation (MC) nor mental-rotation (MR) tasks. Channels placed according to the PROB approach partially overlapped with those from iFMRI and fVASC approaches for MC task. Channels from iFMRI and fVASC approaches overlapped the most, with an average 85.71% [SE = 8.17] for MC and 41.67% [SE = 14.86] for MR. Regarding IS task, P05 showed an overlapping channel between PROB and fVASC layouts (P02 had none). The mean Euclidian distance between the COGs was considerably high (>55mm) for almost all pair of layouts, which indicates that layouts were located in spatially separated areas. IFMRI and fVASC layouts were located, on average, in close proximity for the MC task (6.45mm [SE = 5.64]) and to a lesser extent for MR (42.22 mm [SE = 13.32]). Similarly, the frequency maps shown in the Fig. S9 indicate that (1) the selected channels vary considerably across participants for PROB, iFMRI and fVASC approaches; and (2) iFMRI and fVASC show the highest and most similar spatial extension for MC and MR tasks. As for inner-speech (IS) task, the Euclidean distance ranged between 9.08 mm (PROB-fVASC) and 100.19 mm (LIT-iFMRI) for P05 and between 26.83 mm (LIT-PROB) and 75.98 mm (LIT-iFMRI) for P02 (not shown in Fig. 6).

**Fig 6.**
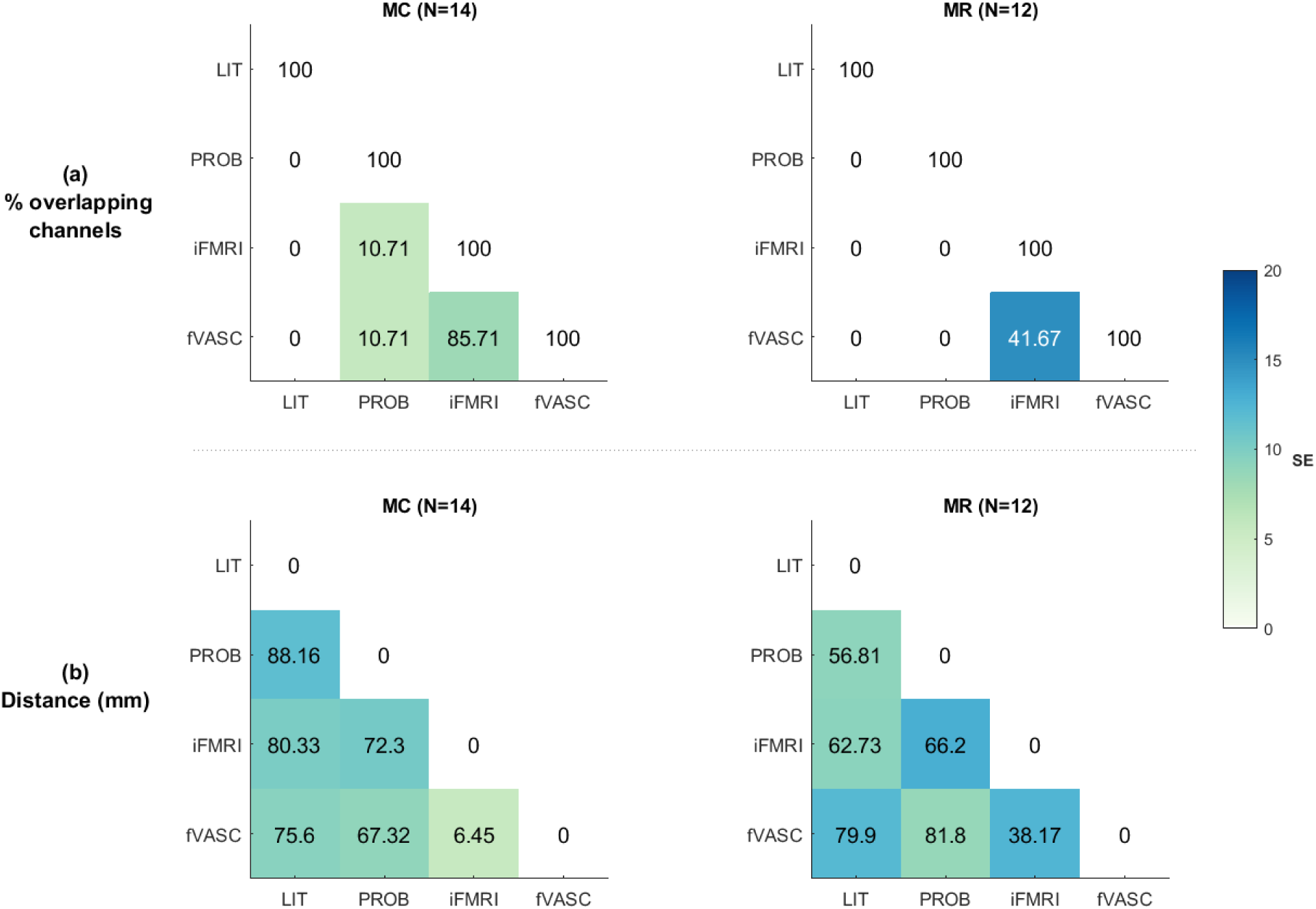
Assessment of degree of layout (dis)similarity across approaches. (a) Average number of overlapping channels for each pair of approach-specific layouts for MC (left) and MR (right) tasks. The numbers in each cell represent the average number of overlapping channels **(a)** or the average Euclidian distance between COG **(b)** for each pair of approach-specific layouts for MC (left) and MR (right) tasks. Colors represent the standard error of the mean. Abbreviations: MC = mental-calculation; MR= mental-rotation.

### 3.2 Significant differences in fNIRS-signal quality across the four optode-placement approaches

The Friedman test was computed separately for each chromophore (Δ[HbO] and Δ[HbR]) and considering (1) all mental-imagery tasks together and (2) each mental-imagery task separately. For Δ[HbO], CNR significantly differed across layouts (Fr = 41.63, df 4,14, p < 0.0001) when all mental imagery tasks were considered together. CNR also differed significantly across layouts for MC (Fr = 24.67, df 3,14 p<0.0001) and MR (Fr = 25.72, df 3,12 p<0.0001). Post-hoc pairwise comparison results with the Wilcoxon signed-rank test and the Benjamini-Hochberg correction method are summarized in Fig. 7. These tests revealed significant differences when all mental-imagery tasks were considered together. Specifically, optodes placed using the LIT approach measured significantly lower CNR values compared to the other three approaches (1) when all mental-imagery tasks were considered together (q[FDR]<0.001), (2) for MC only (q[FDR]_LIT-PROB_ <0.01, q[FDR]_LIT-iFMRI_ <0.001 and q[FDR]_LIT-fVASC_ <0.05) and (3) for MR only (q[FDR] <0.001). In addition, channels placed according to the PROB-derived layout reached significantly lower CNR values than those from the fMRI (q[FDR]_PROB-iFMRI_ <0.001) and fVASC (q[FDR] _PROB-fVASC_ <0.05) approaches.

**Fig 7.**
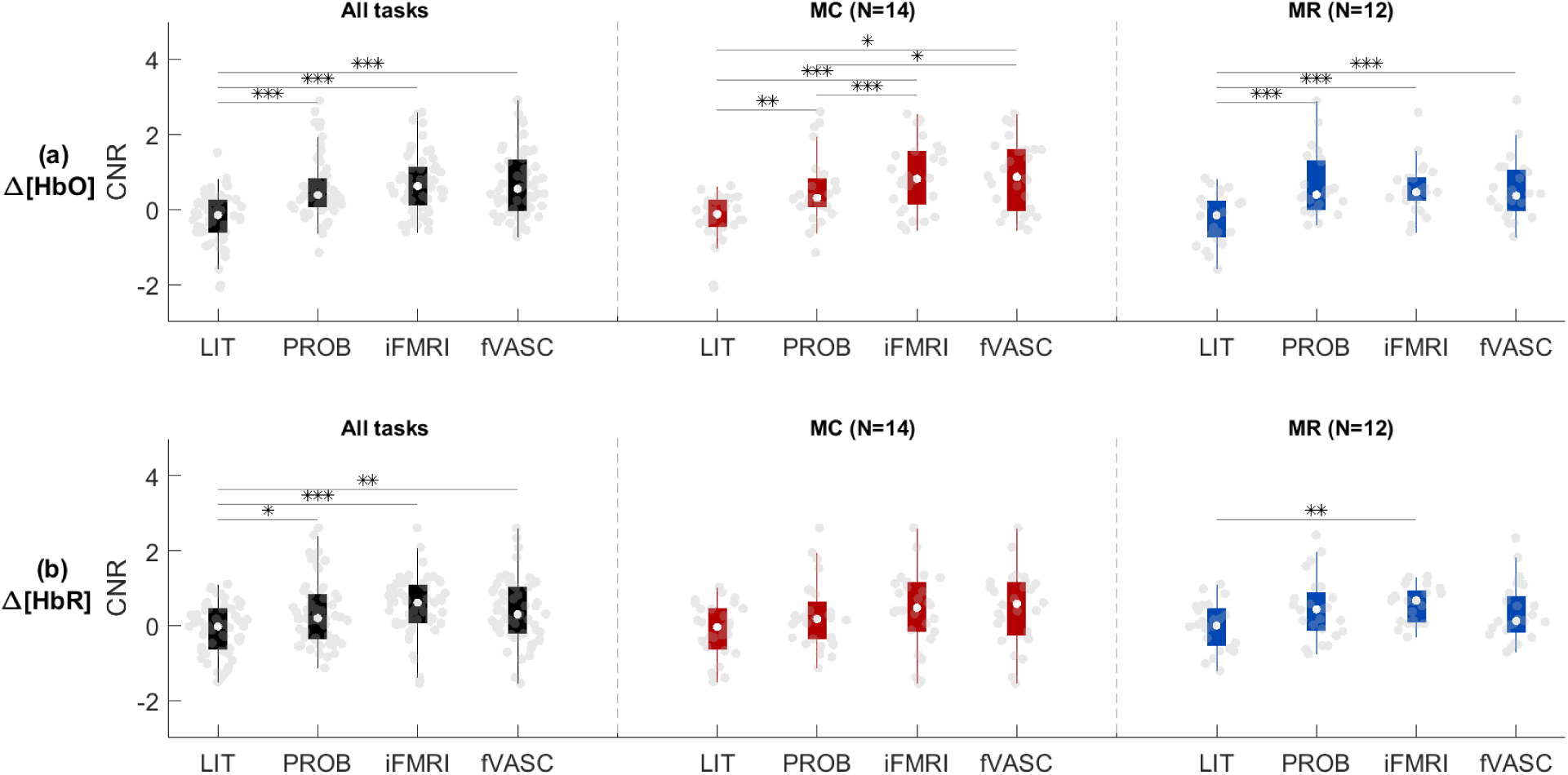
CNR-based group comparison across layouts. Results were evaluated separately for Δ[HbO] (a) and Δ[HbR] (b), when all three mental-imagery tasks were considered together as well as separately for MC and MR tasks (left, middle and right column, respectively). LIT performed significantly worse than the PROB, iFMRI and fVASC approaches for both chromophores when all tasks were considered together. A similar pattern was observed for MC and MR tasks for Δ[HbO]. Gray dots represent single-subject CNR values for a given mental-imagery task. Whiskers represent the 1.5 times the inter-quartile range. Significant parwise differences (calculated using Wilcoxon signed-rank test, one-sided and corrected for multiple comparisons) are indicated with asterisks: *** = q[FDR] < 0.001; ** q[FDR] < 0.01;* q[FDR] < 0.05. Abbreviations: MC = mental-calculation; MR= mental-rotation.

As for Δ[HbR], CNR significantly differed across layouts (Fr = 18.32, df 4,14, p < 0.001) when all mental imagery tasks were considered together. CNR also differed significantly across layouts for MC (Fr = 7.98, df 3,14, p<0.05) and MR (Fr = 8.23, df 3,12, p<0.05). Post-hoc pairwise comparisons revealed that the LIT approach reached significantly lower CNR values when all tasks were considered together for all other layouts (q[FDR]_LIT-PROB_<0.05, q[FDR]_LIT-iFMRI_ <0.001 and q[FDR]_LIT-fVASC_ <0.01). It also reached significantly lower CNR values compared to the iFMRI layout for the MR task (q[FDR]_LIT-iFMRI_ <0.01).

### 3.3 Significant differences in fNIRS-sensitivity across the four optode-placement approaches

#### 3.3.1. T-statistics

For Δ[HbO], ROI *t*-statistics significantly differed across layouts (Fr = 31.66, df 3,14, p < 0.0001) when all mental-imagery tasks were considered together (see Fig. 8). It also differed significantly across layouts for MC (Fr = 23.18, df 3,14 p<0.0001) and MR (Fr = 14.06, df 3,12 p<0.005). *Post-hoc* pairwise comparisons (signed-rank tests, one-sided) revealed that optodes placed using LIT approach measured significantly lower *t*-statistics compared to the other three approaches (1) when all mental-imagery tasks were considered together (q[FDR]<0.001), (2) for MC only (q[FDR]_LIT-PROB_ <0.01 and q[FDR]_LIT-iFMRI; LIT-fVASC_ <0.001) and (3) for MR only (q[FDR] <0.01).

**Fig 8.**
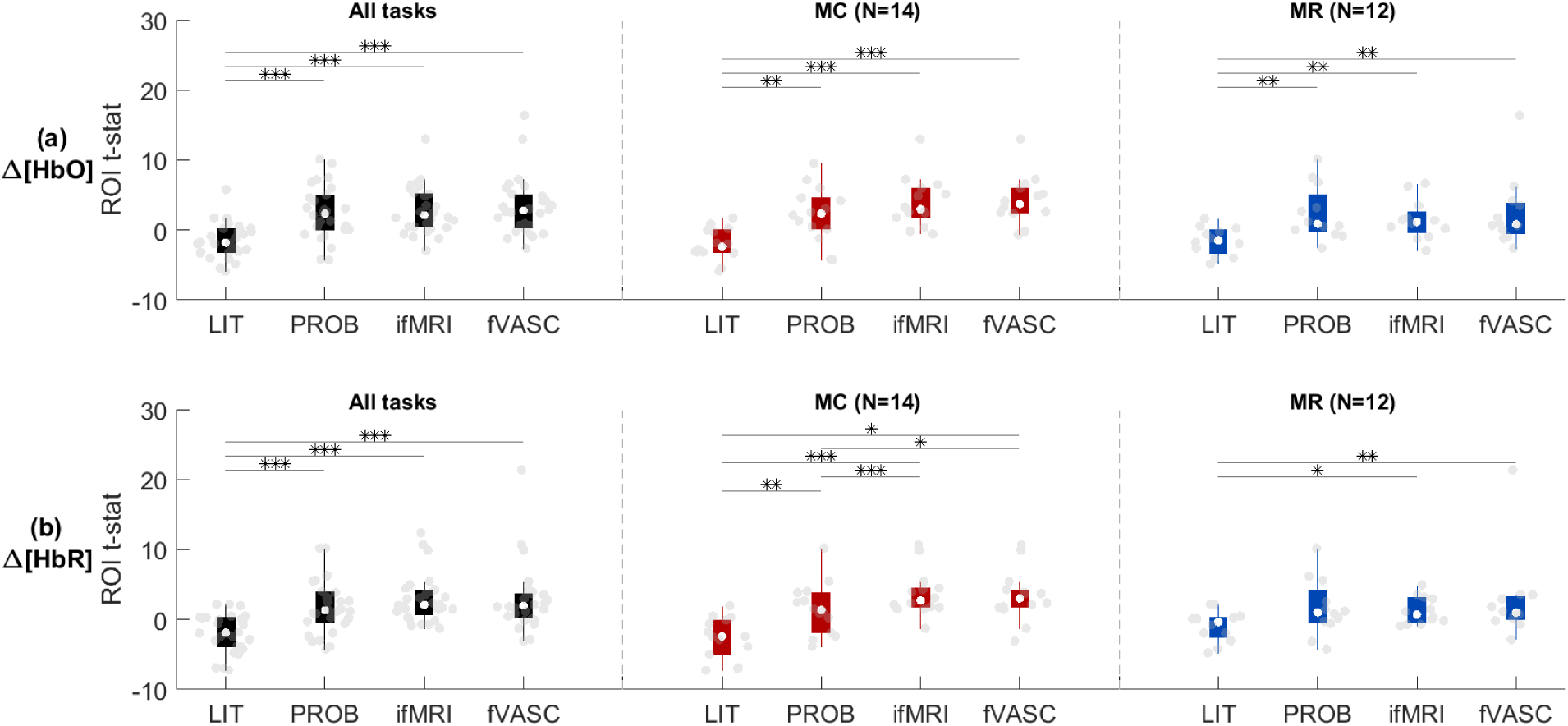
*t-*statistic based group comparison across layouts. Results were evaluated separately for Δ[HbO] (a) and Δ[HbR] (b), when all three mental-imagery tasks were considered together as well as separately for MC and MR tasks (left, middle and right column, respectively). LIT performed significantly worse than the PROB, iFMRI and fVASC approaches for both chromophores when all tasks were considered together. A similar pattern was observed for MC and MR tasks for both chromophores. Gray dots represent single-subject *t-*values for a given mental-imagery task. Whiskers represent the 1.5 times the inter-quartile range. Significant parwise differences (calculated using Wilcoxon signed-rank test, one-sided and corrected for multiple comparisons) are indicated with asterisks: *** = q[FDR] < 0.001; ** q[FDR] < 0.01; *q[FDR] < 0.05. Abbreviations: MC = mental-calculation; MR= mental-rotation.

Δ[HbR] ROI *t*-statistics significantly differed across layouts (Fr = 27.48, df 3,14, p < 0.0001) when all mental imagery tasks were considered together. It also differed significantly across layouts for MC (Fr = 15.46, df 3,14, p<0.01) and MR (Fr = 10.56, df 3,12, p<0.05). *Post-hoc* pairwise comparisons showed a similar trend as HbO: LIT approach measured significantly lower CNR values compared to the other three approaches for almost all comparisons. In addition, the optodes placed according to the PROB approach measured significantly lower CNR than iFMRI (q[FDR]_PROB-iFMRI_ <0.01) and fVASC (q[FDR]_PROB-fVASC_ <0.05).

#### 3.3.2. Percent of participants with significantly active ROIs

Figure 9 shows the percent of participants that resulted in significant activation for each mental-imagery task. For both chromophores, the percent of participants with significant ROI activation increased with increasing the amount of individualized information, and plateaued after including individualized functional maps (for MC task) or was slightly reduced after including vascular information (MR task). For the IS task, PROB and fVASC approaches and PROB and iFMRI approaches contained significant ROI activation for both participants (100%) regarding Δ[HbO] and Δ[HbR], respectively. As for MC and MR tasks, the number of participants with significant activation was higher for more individualized approaches than the LIT approach. Specifically, for the MC task, the LIT approach contained significant ROI activation in 7% (one) participant for both chromophores, while the PROB approach reached significant ROI activation in 57% (eight) and 43% (six) participants for Δ[HbO] and Δ[HbR], respectively. iFMRI and fVASC approaches contained significant ROI activation in 79% (eleven) participants for both chromophores. For the MR task, the LIT approach reached significant activation in 0% and 17% (two) participants, while the PROB approach reached significant activation in 42% (five) and 33% (four) of the participants, while iFMRI and fVASC approaches contained significant ROI activation in 33% (four) and 42% (five) participants for both chromophores, respectively. Figure S10-A shows examples of participants with typical hemodynamic responses (a positive deflection in Δ[HbO] and a negative deflection in Δ[HbR]) for the four approach-specific optode layouts, while Fig. S10-B shows examples of participants with weak/inverted hemodynamic responses for the four approach-specific layouts.

**Fig 9.**
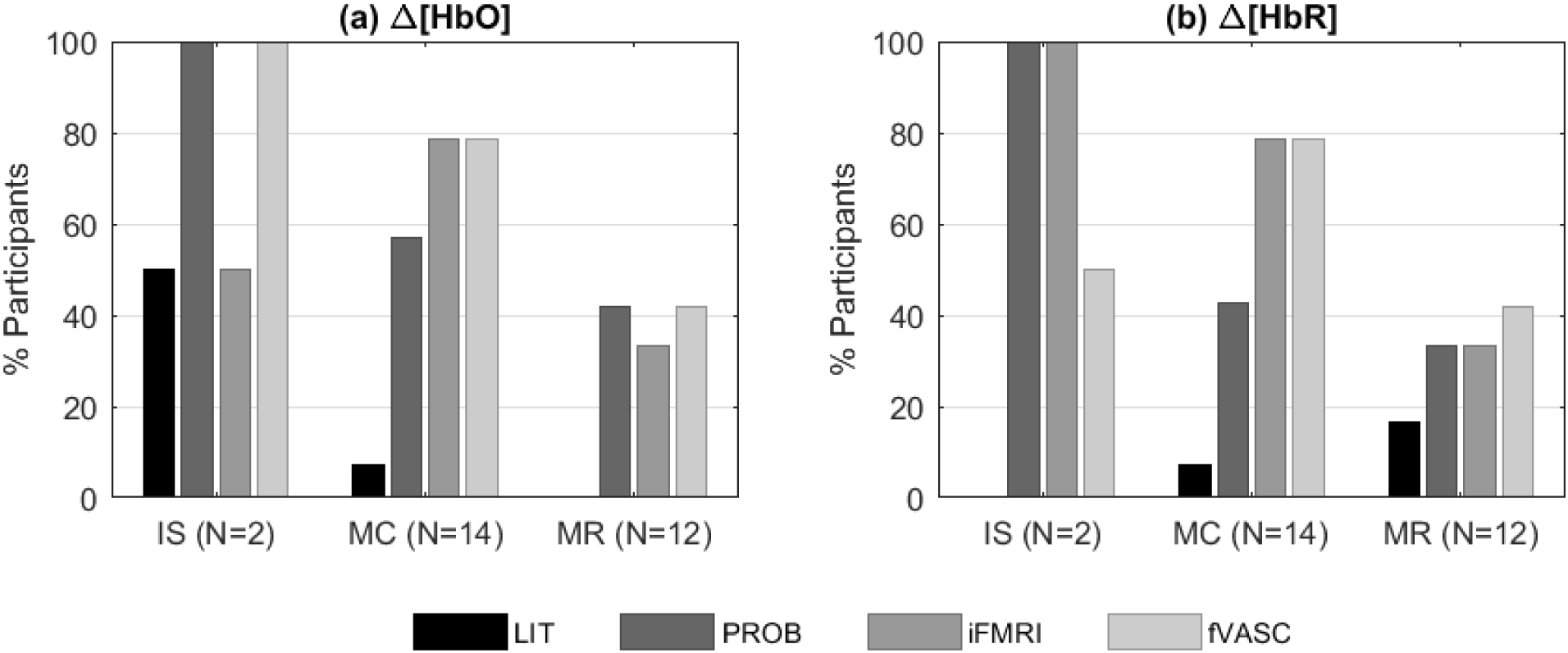
Percent of participants that resulted in significant activation for each mental-imagery task, optode layout and chromophore. For both chromophores, the percent of participants with significant ROI activation increased with increasing the amount of individualized information used to create optode layouts until a certain point: it plateaued after including individualized functional maps (for MC task) or was slightly reduced after including vascular information (MR task). Abbreviations: IS= inner-speech; MC = mental-calculation; MR= mental-rotation.

### 3.4. Spatial specificity of fNIRS-ROIs

To assess how well the fNIRS ROIs targeted individual fMRI activation maps, we computed weighted average and peak fMRI responses within the regions of the cortex interrogated by fNIRS channels. The two plots in Fig. 10a show results for sphere with r=20mm that both the average and peak responses for LIT are significantly lower than the other approaches (significance assessed by signed rank test, one-sided FDR corrected). Using different sphere sizes did not affect the results (data not shown). The temporal correlation between fNIRS and fMRI time courses (bottom plots in Fig. 10b) showed a similar tendency but with smaller differences for Δ[HbO] (and examples of both fNIRS and fMRI time courses are shown in Fig. S11).

**Fig 10.**
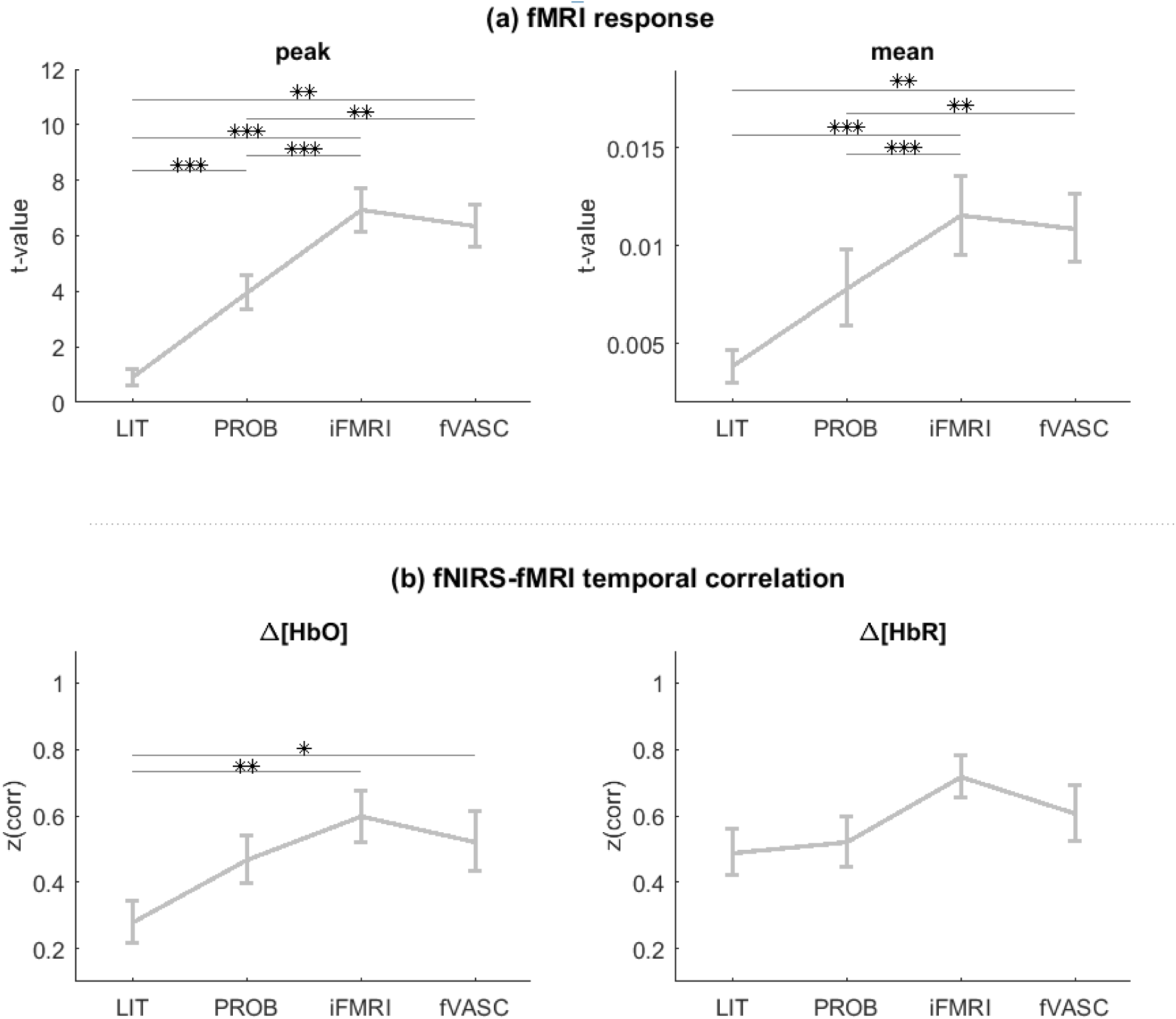
Assessment of layout specificity to fMRI activation maps (a) and of the temporal correlation between fNIRS and fMRI time courses (b). Peak and average values extracted from fMRI activation maps were highest for channels placed according to iFMRI and fVASC approaches and lowest for the LIT approach, independent of the size of projection spheres used to extract the values (data not shown). Times courses of channels placed according to the LIT approach showed significantly lower temporal correlations with fMRI-signal time courses than following the iFMRI and fVASC approaches. Significance was assessed with Wilcoxon paired signed tests (one-tailed) and was corrected for multiple comparisons. *** q[FDR] <0.001; ** q[FDR]<0.01; *q[FDR]<0.05

## 4 Discussion

Designing optode layouts is an essential but challenging step in the preparation of an fNIRS experiment as the quality of the measured signal and the sensitivity to underlying cortex depends on how sources and detectors are arranged on the scalp. This becomes particularly relevant for fNIRS-based BCI and neurofeedback applications, where developing robust systems that use limited number of optodes is crucial to remain practical and comfortable for clinical applications. From the many approaches and tools currently available to optimize optode-layout design, we selected and compared four approaches that incrementally incorporated individual information of participants (LIT, PROB, iFMRI and fVASC) while participants performed mental-imagery tasks typically used in fNIRS-BCI experiments. Our results show that the four approaches resulted in different optode layouts and that the degree of overlap varied across approaches, with the highest overlap and smallest distance between iFMRI and fVASC layouts. Further, time course data of channels placed according to the LIT approach showed significantly lower CNR and *t*-values than those of the channels placed according to the remaining approaches. In addition, we observed no significant difference between PROB, iFMRI and fVASC approaches when all three mental tasks were considered together.

### 4.1 Understanding the difference in performance across layouts

#### Lower performance of the LIT approach

Concurrent fNIRS-fMRI studies show agreement in the hemodynamic signal measured by both modalities (at least in the motor cortex), both temporally ^48^ and spatially ^58^, but inferior in spatial resolution when assessed with fNIRS. To assess how well the fNIRS ROIs targeted individual fMRI activation maps, we computed weighted average and peak fMRI responses within the regions of the cortex interrogated by fNIRS channels. As shown in Fig. 10, the average and peak responses for LIT were significantly lower than the remaining approaches. The temporal correlation between fNIRS and fMRI time courses showed a similar tendency. These observations were expected since PROB, iFMRI and fVASC approaches were based on fMRI information. However, if the individual fMRI map is used as the ground-truth measure of cerebral activity due to its superior resolution and higher SNR, Fig. 10 shows that the LIT approach could not capture the underlying signal as good as the other approaches.

Several factors may have contributed to that. First, the head model used for Monte Carlo simulations for LIT differed from the other three approaches (Colin27 head atlas *vs.* subject-specific anatomical model, respectively). Although head atlases are good approximations, the tissue geometries may significantly differ from other adult individuals ^39^. Second, the ROI selection procedure for LIT differed from the PROB, iFMRI and fVASC approaches in that the ROI selection for LIT was based on a literature review, while the other three approaches relied on functional contrast maps. Due to the small number of participants for the IS task (N=2), the following lines will focus only on MC and MR tasks. The mental-imagery instructions used in this study differed from the reviewed studies, which may have contributed to a suboptimal selection of the ROIs for the LIT approach. Indeed, the majority of reviewed papers that reported using mental arithmetic used strategies that aimed at increasing the working memory demand and thus mainly measured brain activation in the frontal lobe. Examples of tasks used in these studies are subtraction to visually presented 3-digit or 2-digit numbers, or addition or multiplication of visually presented single or double digits to/with single or double digits ^13, 14, 59–72^. Here, we asked participants to recite common multiplication tables, which is considered an easy task and thus may have elicited lower responses in frontal and parietal areas when compared to more complex multiplication problems ^73^. Regarding mental rotation, most of the reviewed work used visually presented cues that had to be mentally rotated, such as geometric object, alphanumeric character or hand rotations ^61, 74–84^. In this study, we did not visually present the object to be mentally rotated, as participants were asked to imagine a diver spinning in the air while keeping their eyes closed. In addition, unlike the reported studies, there was no reference object to compare to the rotated object. The lack of visual support and a reference object could cause the recruitment of the areas involved in the task to be slightly different or to be recruited to a lesser extent. However, we would like to note that since we did not test the performance of approaches that use anatomical ROIs defined on individual head models, we cannot disentangle if the lower performance of the LIT approach is due to the head model used or whether it is due to the nature of the ROI.

#### No significant difference between fVASC and iFMRI layouts

The fVASC and the iFMRI approaches only differed in the number of tissues used during Monte Carlo simulations: the fVASC condition included an additional participant-specific vascular information. Including an additional vascular information did not result in a significant difference compared to the iFMRI layout at the group level. This is mainly because the generated layouts were very similar between them, as indicated by the channel overlap across layouts and the Euclidian distance (Fig. 6). This high similarity seems to be driven by the functional ROIs, which was the same for both approaches. Our decision to use a small number of optodes for each layout, the constraints to select them, and segmentation-related factors (see the limitations section 4.2) may have also limited the improvements expected from the fVASC approach.

#### PROB performs similar to iFMRI and fVASC

Here we defined probabilistic functional activation maps for each participant and task from an independent dataset using a leave-one-subject-out scheme. This approach resulted in an improved sensitivity that is comparable with the improvement observed when using individual data of a given participant. Specifically, we observed that CNR and *t*-statistics performed similarly for the PROB approach compared to the iFMRI and fVASC approaches. Further, Figure 10 also shows that, descriptively speaking, the peak and average values captured by channels defined based on the PROB approach are closer to those of iFMRI and fVASC approaches than the LIT approach is. This is because PROB approach-based activation maps show high spatial correspondence when compared to the reference fMRI maps for each participant and mental task. Indeed, the average spatial correlation (assessed by Spearman correlation) between probabilistic maps and individual activation was 0.63 when all tasks are considered together and of 0.63 for MC and 0.64 for MR tasks. For IS the values ranged between 0.52 and 0.66. These values, together with the results presented in this study, suggest that using probabilistic maps based on a reasonable number of participants and defined on individual anatomical space can be used for any new participant (as long as the functional maps used to create the probabilistic maps are based on the same task or are closely related to it).

### 4.2 Optode-layout design and its limitations

#### Cost function, constraints and optimization problem

The optode layout for each of the four approaches consisted of two channels that shared one optode and that maximized the total sensitivity to the preselected ROI. The cost function to be maximized was the same as in ^5^, but the algorithmic approach to solve the optimization problem was tailored to account for the constraints imposed by our particular research question(s) and experimental design. This entails that our algorithmic approach may not be (and was not designed to be) generalizable to other experimental designs. Importantly, although the approach by ^5^ is very effective in covering focal ROIs, it fails to provide an appropriate solution when the ROI is extended or consists of multiple noncontiguous regions ^1, 5^, as was the case in this study.

#### Mental-imagery task selection

From the three mental-imagery tasks participants had to perform inside the MRI scanner, each participant performed two tasks during the fNIRS session. The selection of these two tasks was subject-specific and followed several pre-defined criteria. Combining approach-specific layouts for both mental-imagery tasks caused incompatible source and detector placements in some participants. The decisions taken to overcome these problems, together with the subject-specific task selection led to an unequal number of participants for each task (N_IS_ = 2, N_MC_ = 14, N_MR_ =12), which made the group analysis for IS task unfeasible. To overcome the incompatibility problem, future studies could test the performance of different layouts in different runs/sessions (by using a given layout at a time), whose order could be counter-balanced to account for run/session effects. In addition, a single mental-imagery task could be studied at a time (instead of multiple tasks as in this study).

#### Monte Carlo simulations

Our light sensitivity profiles may contain estimation errors due to a number of simplifications. First, the head models used in this study did not consider that the skull can contain cancellous and cortical bone, and the soft tissue may contain fat and muscle that have different optical properties^85^. Second, both sources and detectors were modeled as pencil sources instead of separately being modelled according to their function (they emit or detect light) and technical characteristics. Third, we did not distinguish between arteries and veins when defining the head model. Even if our decision can be justified by the relatively small difference in optical properties between veins and arteries compared to the remaining tissues, we cannot discard potential divergence in the results if arteries and veins had been distinguished. Optical properties also differ depending on the diameter of blood vessels ^86^, which we did not take into account in the current study. Finally, our vascular maps depended on manual segmentation procedures, which may have introduced variability. Future studies may overcome these limitations by mapping superficial (scalp/skull) vasculature with more optimized MRI sequences ^87^, and by distinguishing between arteries and veins and their diameters ^88, 89^.

### 4.3. Implications for BCI applications

In fNIRS-BCI applications for motor-independent communication and control, brain responses from a set of tasks are discriminated by exploiting information in distributed patterns of brain activity using multi-channel pattern analysis (the equivalent to multi-voxel pattern analysis in fMRI studies). Alternatively, univariate analysis in combination with temporal encoding paradigms can be used, where participants perform a (number of) task(s) in a specific time window and evoke brain activation in a single (distinct) brain location(s) ^10–12, 16, 90, 91^. For either approach, it is important to ensure there is a set of channels that contains sufficient task-related information to discriminate responses.

The present study constitutes a relevant pre-step for these BCI applications as it compared approaches that used different amount of individualized information to design task-specific, optimized optode layouts that should result in informative channels. Neurofeedback applications can also benefit from layouts that ensure sufficient task-related information and improved spatial specificity. Our results show that the percent of participants with significant ROI activation increased when increasing the amount of individualized information to create the optode setup, but only until a certain point. Indeed, adding vasculature information did not increase the percent of participants for MC and reduced this number for MR. Although all participants showed significant activation levels for every mental task during the fMRI run, none of the approaches using fMRI information managed to get all participants to have significant ROIs for MC and MR tasks. It is unclear whether a given level of fMRI activation is enough to guarantee the detection of task-related fNIRS signal. Even if both neuroimaging methods measure the hemodynamic response to neural activity, fNIRS is highly dependent on the individual anatomical features, such as the scalp-brain distance (which differs across the head)), the presence of hair, etc.^18^. In addition, our fNIRS results might have been affected by the discrete spatial locations used in this study (130 EEG positions). Spatially unrestricted optode placement would likely improve the results substantially^92^.

### 4.4 Recommendations for optode placement and the way forward

Effective optode-layout design balances a number of potential tradeoffs. The extended layouts based on the international 10-20 system or its extensions can be used to study functional network dynamics and are adequate when target ROIs are not easy to define ^92^. In addition, although the target ROI may not be optimally sampled (due to unavoidable regions not covered by a source-detector pair when creating optode layouts and the lower spatial resolution associated to fNIRS compared to fMRI), the chance of completely missing it is relatively low. That said, smaller setups are preferred in fNIRS-BCI applications due to their superior practicability and patient comfort. However, they run a much higher risk of missing signal from the target ROI due to anatomical or functional differences between individuals. As a result, small BCI setups are likely to benefit from supplementary f/MRI data investigated in the present work. The recommendations and conclusions presented here therefore focus on this particular fNIRS application.

Considering that any additional individualized information has an associated acquisition/analysis cost, it is worth asking, especially when temporal/monetary/material resources are limited: how much individual information is worth to include for designing optode layouts? Figure 11 shows the predicted percent improvement in performance (in terms of *t*-statistics [top] and CNR [bottom]) *vs.* the additional time required to acquire/analyze the data relative to the LIT layout, here considered the “baseline” approach. Points above the line indicate that the percent improvement of a given performance measure is higher than the temporal resources spent to achieve that gain. The figure suggests that including individual anatomical data (PROB layout) or including both, individual anatomical and functional data (iFMRI layout), improves the performance while efficiently using temporal resources. It also suggests that the fVASC approach in its current form is not as cost-effective.

**Fig 11.**
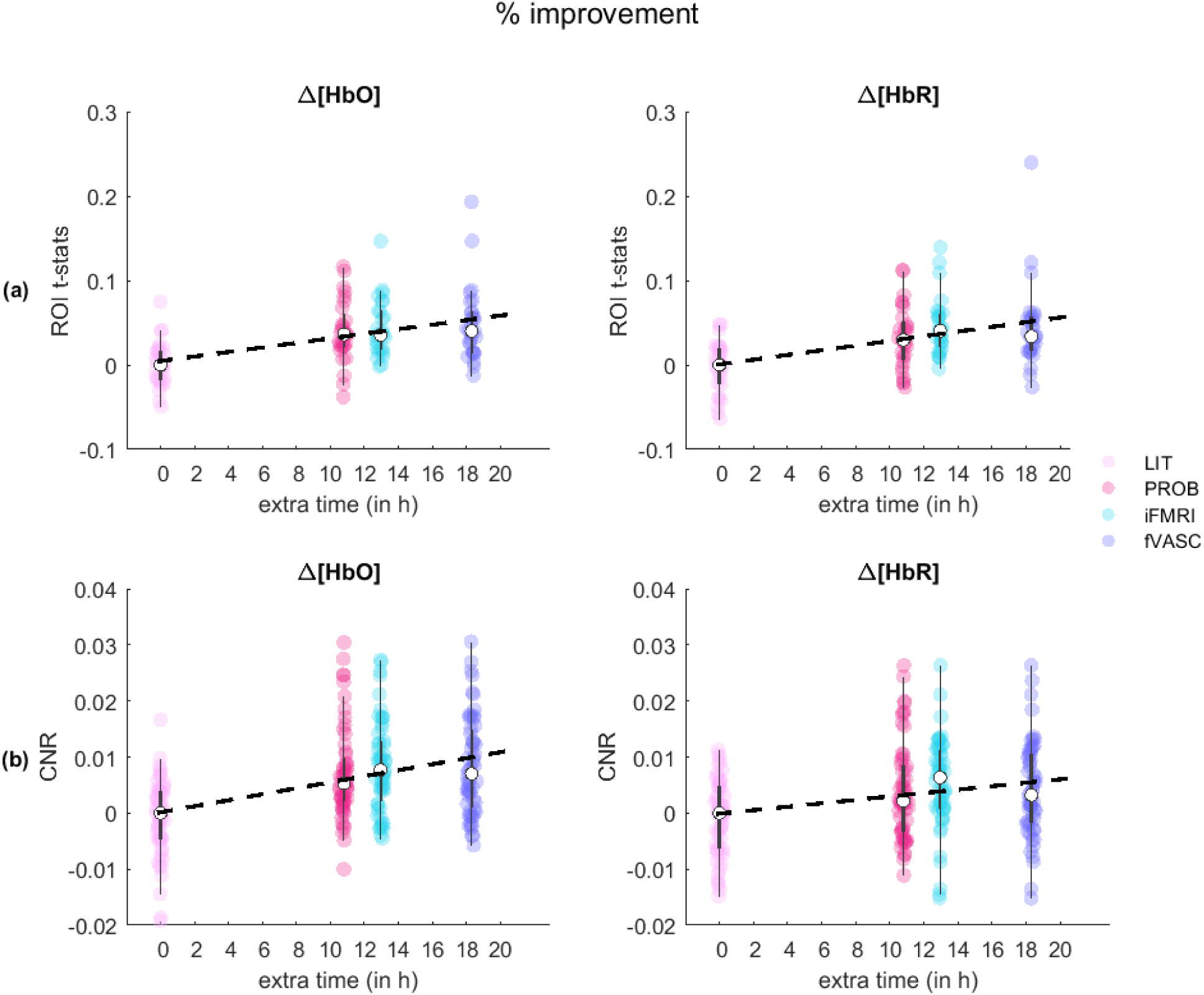
Percent improvement in performance (in terms of *t*-statistics (a) and CNR (b)) *vs.* the additional time required to acquire/analyze the data (in hours). All values are relative to the LIT approach (in light pink), here considered the “baseline”. The bigger white circles represent the median of the percent improvement in *t*-statistics/CNR values for each layout when all three tasks are considered together. The dashed line represents the predicted percent improvement in performance for a given processing time. Points above/below the line indicate that the percent improvement of a given performance measure is higher/lower than the temporal resources spent to achieve that gain.

The analysis described above focused only on a small part of the multi-dimensional problem related to cost-effectiveness. Naturally, costs and benefits of including more individualized information for creating clinically practical layouts should be assessed in that very context. For example, in certain (rare) cases such as long-term BCIs in ‘locked-in’ patients, using individual (f)MRI data may result in increased ability to communicate, i.e., provide considerable benefit. In that case, even though using individual (f)MRI is more resource-demanding, the benefits could outweigh the costs.

In view of these observations, we encourage researchers to use individual functional and anatomical data for designing optode layouts when possible, but when anatomical data is available and functional data is not, probabilistic functional maps constitute a promising and economic alternative. FMRI-based probabilistic functional maps of the human ventral occipital cortex ^93^, human motion complex ^94^, face selective areas ^95, 96^, finger dominance in the primary somatosensory cortex ^97^ or across the whole cortex ^36^ are freely available or available on demand. However, we could not find any published work on probabilistic mental-imagery maps, which could be beneficial for optode placement in BCI research. To improve this situation, the probabilistic functional maps of the three mental-imagery tasks used in this study (in MNI space) are available upon request. Finally, in the absence of functional and anatomical information, ROI selection should be guided by relevant body of work or meta-analyses that describe tasks closely related to the ones to be used during the fNIRS session. In parallel, a larger setup could be initially employed in a “localizer” run to determine the most informative channels which could be subsequently scaled down to consider only the most informative channels. In the present study, once the target ROIs were selected, we used FOLD ^30^ for designing our optode layout due to its user-friendly features. However, other toolboxes such as Array Designer ^1^ and software, such as NIRStorm (a *BrainStorm* plugin for fNIRS analysis ^28^), also offer promising and flexible tools that were not explored in the present study.

## 5 Conclusions

In this paper, we compared four approaches to design small fNIRS optode layouts that represent various scenarios research groups may encounter when planning fNIRS-BCI experiments. By providing the insights of such comparisons, we hope to have offered an informative framework so that researchers can efficiently use resources for developing robust and convenient fNIRS-BCI systems for clinical use.

## 7 Acknowledgements

The authors would like to thank Agustin Lage-Castellanos, Andrew Morgan, Faruk Gülban and Judith Eck for their advice on data analysis; and to all the participants for their time. The authors would also like to thank NIRx for providing the short-distance optode bundles.

This work was supported by the European Commission (7th Framework Program 2007-2013, DECODER project, to BS and RG), by the Netherlands Organization for Scientific Research (Research talent grant 406-15-217 to LN-C) and by The Luik 3 grant for joint research between Cognitive Neuroscience and Knowledge Engineering Departments, Maastricht University, on advanced brain-robot interfaces, 2015-2019 (to RM, RG and BS).

## Author Biographies

**Amaia Benitez-Andonegui** received a BS in Biomedical Engineering at Tecnun School of Engineering and a MS in Cognitive and Clinical Neuroscience at Maastricht University. She is currently a PhD candidate at the Cognitive Neuroscience and Knowledge Engineering departments at Maastricht University. Her research interests focus on advancing the applicability of fNIRS-based brain-computer interface applications in healthy and clinical populations.

**Michael Lührs** is a postdoctoral researcher in the department of Cognitive Neuroscience at Maastricht University and BrainInnovation BV. His main research focus is the advancement of neurofeedback tools and other real-time applications using fMRI and fNIRS.

**Laurien Nagels-Coune** received a BS and MS in Psychology from the University of Leuven and a MS in Cognitive and Clinical Neuroscience from Maastricht University. Currently, she works as a PhD student at the Department of Cognitive Neuroscience at Maastricht University. Laurien does research in translational neuroscience using fNIRS-based brain computer interfaces for communication with locked-in patients. Laurien also works as a neuropsychologist in a division for noncongenital brain damage in Belgium.

**Dimo Ivanov** is an Assistant Professor in the MR Methods group at the Department of Cognitive Neuroscience and Maastricht Brain Imaging Centre. His research interests range from anatomical imaging through perfusion mapping to high-resolution functional MRI.

**Rainer Goebel** is a full professor in the Department of Cognitive Neuroscience at Maastricht University. He received several grants from the Human Brain Project (2014-2023). He founded the company Brain Innovation BV that produces free and commercial software for neuroimaging data analysis and clinical applications. He was selected as member of the Royal Netherlands Academy of Arts and Sciences (2014) and Leopoldina, the German National Academy of Science (2017).

**Bettina Sorger** is an Associate Professor at the Department of Cognitive Neuroscience Maastricht Brain Imaging Centre. Her research interests include motor-independent communication/control and neurofeedback (therapy) - both based on brain hemodynamics.

