## Supplementary Figures and Tables for "Guiding functional near-infrared spectroscopy optode-layout design using individual (f)MRI data: Effects on signal quality and sensitivity"

### A. Materials and Methods

#### *A.1 Preprocessing and analysis of functional MRI data*

<u>fNIRS coverage mask definition</u>

A whole-head mask was created for each participant from the bias-corrected structural image. Each mask was iteratively eroded 50 times and all voxels that did not belong to this eroded mask were selected and intersected with the original head mask. The 'surviving' voxels were used to mask out active voxels from deeper regions, as we did not expect the fNIRS signal to be sensitive to these regions (Strangman et al., 2013).

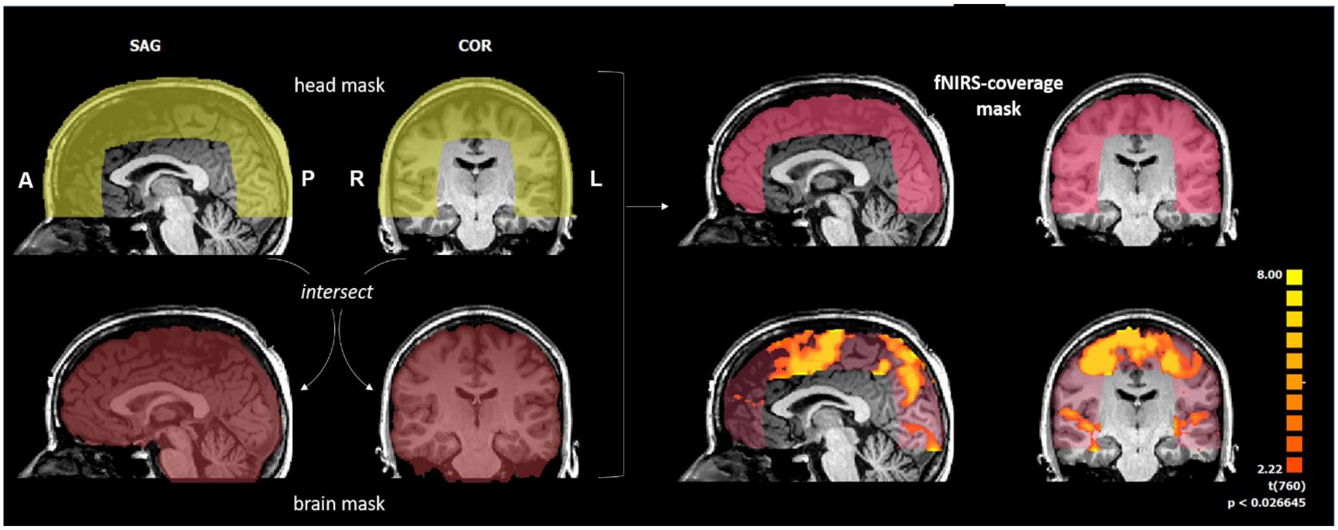


***Fig. S1. fNIRS-coverage mask definition and application**. An fNIRS-coverage mask was created by the intersection of the eroded head mask and the brain mask, and it was used to mask out the active voxels from deeper regions. The activation map depicted in this figure resulted from the MR vs. Rest contrast for a representative participant. The activation map was corrected using a cluster-extent threshold at 5%.*

<u>Probabilistic functional maps</u>

We defined subject-specific probabilistic functional maps based on an independent sample, i.e., the functional data from the remaining individuals. Fig. S2 depicts an example of probabilistic functional maps for each of the mental-imagery task, from a left, top and right view. Colors represent the percent overlap of significant activation across participants for a task vs. rest contrast (report correction)

Vascular segmentations and reconstructions

Figure S3 summarizes the steps carried out for segmenting cerebral, pial and scalp vessels and Fig. S4 depicts the resulting vascular reconstruction for a sample participant. Cerebral/pial vessels are shown in blue, while scalp vessels are shown in red. Scalp vasculature segmentation required more manual corrections than the cerebral/pial vasculature segmentation.

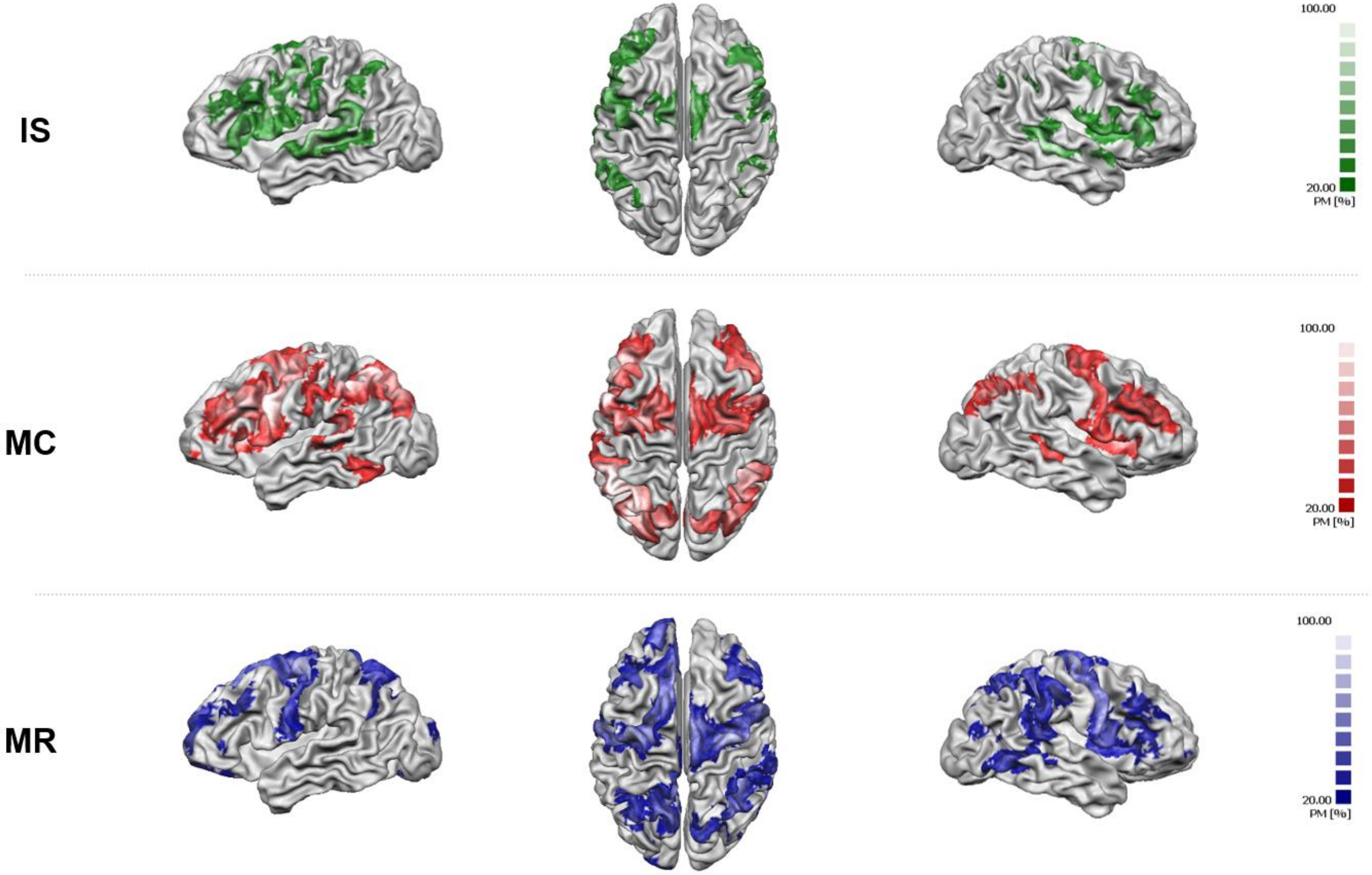


***Fig. S2. Example of functional probabilistic maps (left, top and right view) for each mental-imagery task (IS= inner speech; MC = mental calculation; MR = mental rotation).***

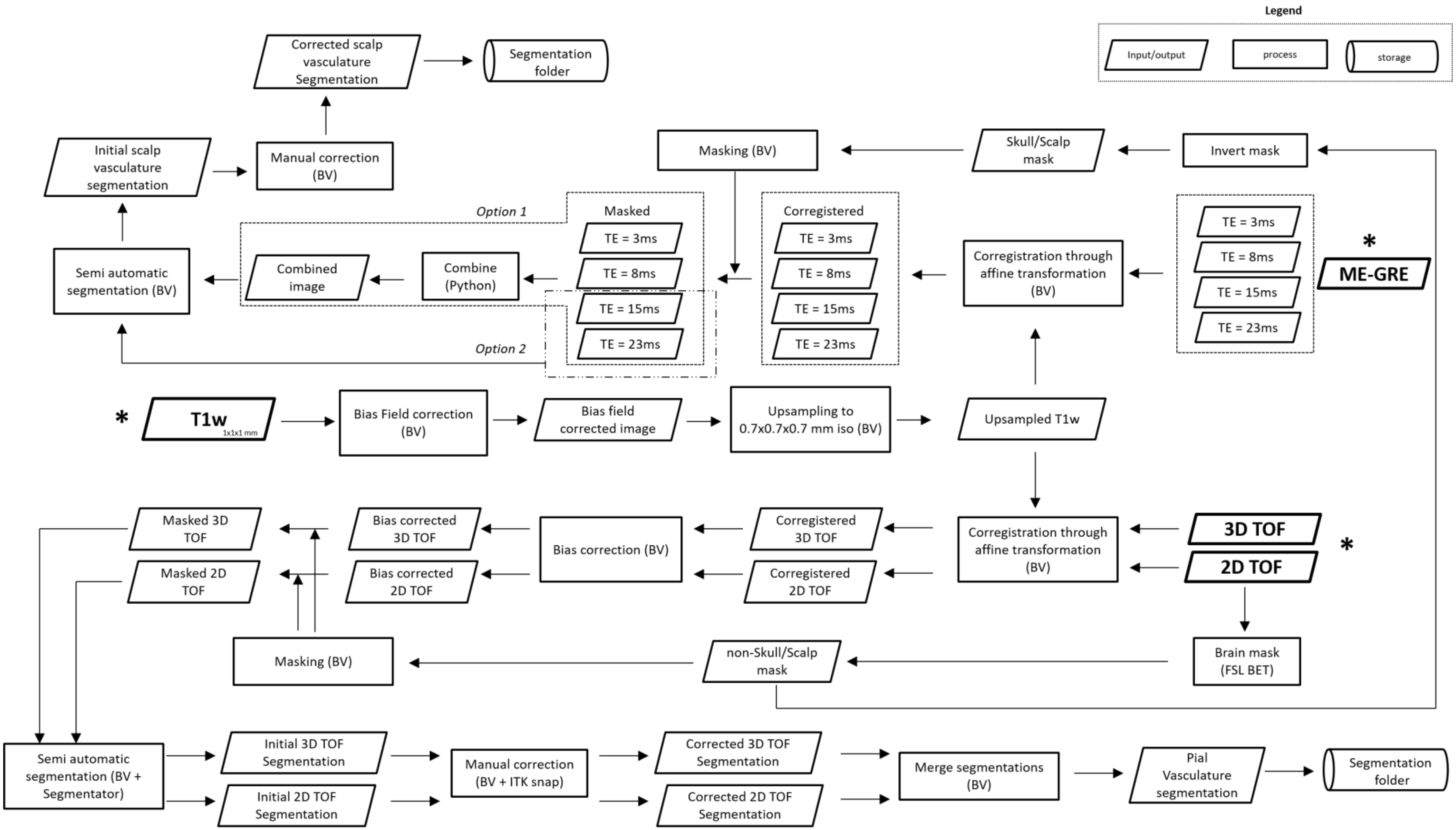


***Fig. S3. Steps involved in vascular data segmentation for each participant.*** *Asterisks indicate starting points in the pipeline. BV = Brainvoyager QX; TOF = time-of-flight; T1w = structural (MPRAGE) image; ME-GRE: multi-echo gradient echo images.*

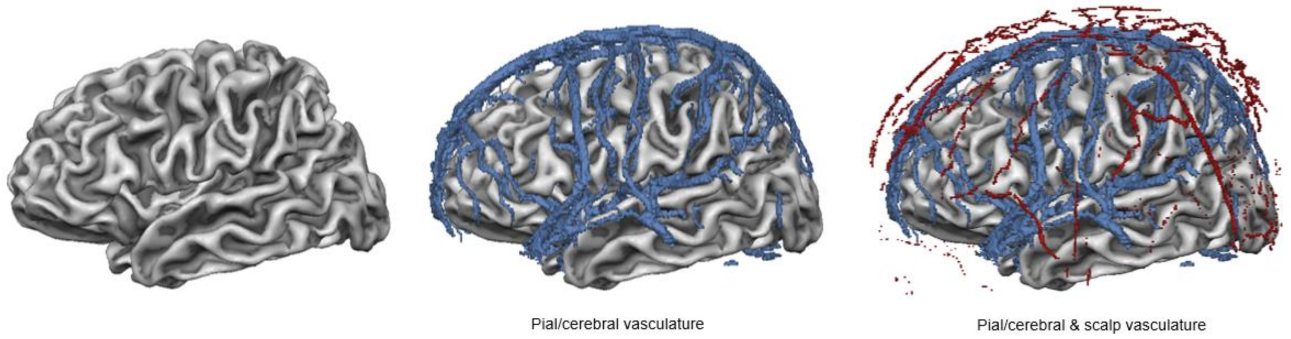


***Fig. S4. Example of vascular reconstructions (P16).*** *Left. Cortical reconstruction (left hemisphere). Middle. Pial/cerebral vasculature reconstruction (in blue) overlaid onto the cortical mesh. Right. Scalp vasculature reconstruction (in red) overlaid onto the cortical mesh and the pial/cerebral vasculature reconstruction.*

#### *A.2 Optode layout creation*

<u>Monte Carlo simulations</u>

*Head models*

Figure S5 shows the head models used for PROB (left figure), iFMRI (left figure) and fVASC (right figure) during Monte Carlo simulations. Both figures differ in the amount of tissues included: while the left figure represents a five-tissue model, the right figure includes a sixth tissue (vascular structures). Importantly, we did not distinguish between arteries and veins.

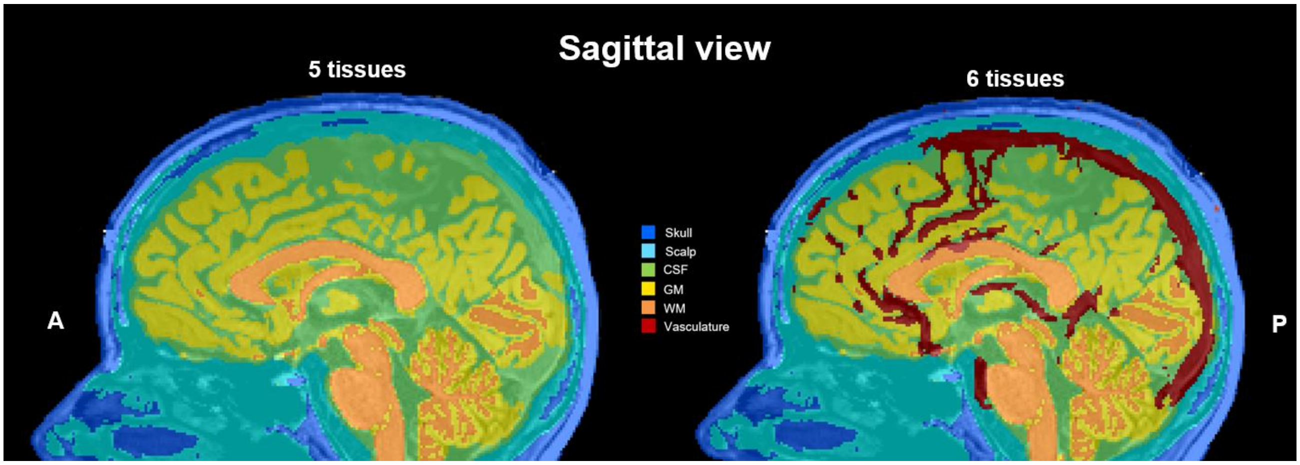


***Fig. S5*. Example head model (P01) used for Monte Carlo simulations.** (Left) Five-tissue model used for PROB and iFMRI. (Right) Six-tissue model used for fVASC.

*Optical properties for vascular structures*

The oxygen saturation of blood ($SO_2$) is defined as the ratio of the $HbO_2$ concentration to the total hemoglobin concentration, where $SO_2$[Arteries] ~ 97.5% and $SO_2$[Veins] ~ 75%. Bosschaart et al.

(2014) reported optical coefficients for a range of wavelengths and $SO_2$ of 98% and $SO_2$ of 0%. $SO_2$=98% can be considered an approximation of the arterial blood and a combination of both ($SO_2$=98% and $SO_2$=0%) can be used to approximate the venous blood:

$$SO_2[Veins] = 0.75 \times SO_2[98\%] + 0.25 \times SO_2[0\%] \quad (1)$$

***Table S1. Optical values reported in Bosschaart et al. (2014) for the four wavelengths used in the present study.*** *The scattering (μs) and anisotropy (g) parameters were based on a mathematical model (proposed model, column 1 in the present table), while the absorption parameters (μa) based on empirical data (column 2). The columns in red correspond to optical values for venous blood which was calculated based on a weighted sum of SO2 98% and 0% values (equation 1).*

| | proposed model | | | | | | empirical data | | |
|---|---|---|---|---|---|---|---|---|---|
| **λ (nm)** | **μs ($mm^{-1}$) SO2 (98%)** | **μs ($mm^{-1}$) SO2 (0%)** | **μs SO2 (75%)** | **g SO2 (98%)** | **g SO2 (0%)** | **g SO2 (75%)** | **μa ($mm^{-1}$) SO2 (98%)** | **μa ($mm^{-1}$) SO2 (0%)** | **μa SO2 (75%)** |
| 690 | 85,84 | 75,63 | 83,2875 | 0,9843 | 0,9852 | 0,9845 | 0,13 | 1,17 | 0,39 |
| 750 | 77,04 | 67,62 | 74,685 | 0,9827 | 0,9836 | 0,9829 | 0,24 | 0,81 | 0,3825 |
| 780 | 72,59 | 63,73 | 70,375 | 0,9819 | 0,9827 | 0,9821 | 0,33 | 0,59 | 0,395 |
| 830 | 66,96 | 58,72 | 64,9 | 0,9804 | 0,9812 | 0,9806 | 0,46 | 0,43 | 0,4525 |

***Table S2. Optical properties of arterial and venous blood based on the optical values reported in Bosschaart et al. (2014).*** *Values were computed as the average of four wavelengths used in the present study.*

| | **μa ($mm^{-1}$)** | **μs ($mm^{-1}$)** | **g** |
|---|---|---|---|
| **Arterial blood (SO2 98%)** | 0,29 | 75,6075 | 0,982325 |
| **Venous blood (SO2 75%)** | 0,405 | 73,311875 | 0,982525 |

<u>LIT approach: ROI definition</u>

*Studies included in the literature review*

We used the PubMed database and keywords for the search included: ‘inner speech’, ‘covert speech’, ‘mental talking’ and ‘overt speech’ for inner speech (overt speech was also included as both covert and overt speech share common networks, see, *e.g.*, Martin et al. (2014)); ‘mental calculation’ and ‘mental arithmetic’ for mental calculation; and ‘mental rotation’ for mental rotation. We included every work that reported using such mental-imagery tasks, independent of the neuroimaging modality and the population under study (see Table S3).

*Selected ROIs*

ROIs for each mental-imagery task were selected based on the most-frequent regions reported across studies. This number different across the three tasks (see Table S4).

76 ***Table S3. Literature review used to select ROIs for the LIT approach.***

| Author | Title paper | Task | | | Method |
|---|---|---|---|---|---|
| Shergill et al. (2001) | A functional study of auditory verbal imagery | IS | | | fMRI |
| Baciu et al. (1999) | fMRI assessment of hemispheric language dominance using a simple inner speech paradigm | IS | | | fMRI |
| Fujimaki et al. (2004) | Right-lateralized neural activity during inner speech repeated by cues | IS | | | fMRI + MEG |
| Hurlburt et al. (2016) | Exploring the Ecological Validity of Thinking on Demand: Neural Correlates of Elicited vs. Spontaneously Occurring Inner Speech | IS | | | fMRI |
| Girbau (2014) | A Neurocognitive Approach to the Study of Private Speech | IS | | | fMRI, PET, ERP, MEG |
| Cannestra et al. (2003) | Functional assessment of Broca's area using near infrared spectroscopy in humans | IS | | | fNIRS |
| Wan et al. (2018) | A functional near-infrared spectroscopic investigation of speech production during reading | IS | | | fNIRS |
| Zhang et al. (2017) | Signal processing of functional NIRS data acquired during overt speaking | IS | | | fNIRS |
| Aziz-Zadeh et al. (2005) | Covert Speech Arrest Induced by rTMS over both Motor and Nonmotor Left Hemisphere Frontal Sites | IS | | | rTMS |
| Martin et al. (2016) | Word pair classification during imagined speech using direct brain recordings | IS | | | ECOG |
| Yoo et al. (2004) | Brain computer interface using fMRI: spatial navigation by thoughts | IS | MC | | fMRI |
| Sereshkeh et al. (2018) | Online classification of imagined speech using functional near-infrared spectroscopy signals | IS | | | fNIRS + EEG |
| Herff et al. (2012) | Cross-Subject Classification of Speaking Modes Using fNIRS | IS | | | fNIRS |
| Morin and Michaud (2007) | Self-awareness and the left inferior frontal gyrus: Inner speech use during self-related processing | IS | | | fMRI [review] |
| Verner et al. (2013) | Cortical oxygen consumption in mental arithmetic as a function of task difficulty: a near-infrarred spectroscopy approach | | MC | | fNIRS |
| Pfurtscheller et al. (2010) | Focal frontal (de)oxygemoglobin responses during simple arithmetic | | MC | | fNIRS |
| Kawashima et al. (2004) | A functional MRI study of simple arithmetic - a comparison between children and adults | | MC | | fMRI |
| Rickard et al. (2000) | The calculating brain: an fMRI study | | MC | | fMRI |
| Power et al. (2012) | Automatic single-trial discrimination of mental arithmetic, mental singing and the no-control state from prefrontal activity: toward a three-state NIRS-BCI | | MC | | fNIRS |
| Shin et al. (2016) | Near-infrared spectroscopy (NIRS)-based eyes-closed brain-computer interface (BCI) using prefrontal cortex activation due to mental arithmetic | | MC | | fNIRS |
| Naito et al. (2007) | A Communication Means for Totally Locked-in ALS Patients Based on Changes in Cerebral Blood Volume Measured with Near-Infrared Light | | MC | | fNIRS |
| Weyand and Chau (2015) | Correlates of Near-Infrared Spectroscopy Brain–Computer Interface Accuracy in a Multi-Class Personalization Framework | WG | MC | | fNIRS |
| Bauernfeind et al. (2008) | Development, set-up and first results for a one-channel near-infrared spectroscopy system | | MC | | fNIRS |
| Ogata et al. (2007) | A Study on the Frontal Cortex in Cognitive Tasks using Near-Infrared Spectroscopy | WG | MC | | fNIRS |
| Utsugi et al. (2007) | Development of an Optical Brain-machine Interface | WG | MC | | fNIRS |

| | | | | | |
|---|---|---|---|---|---|
| Ang et al. (2012) | Extracting and selecting discriminative features from high density NIRS-based BCI for numerical cognition | | MC | | fNIRS |
| Schudlo et al. (2013) | Dynamic topographical pattern classification of multichannel prefrontal NIRS signals | | MC | | fNIRS |
| Schudlo and Chau (2013) | Dynamic topographical pattern classification of multichannel prefrontal NIRS signals: II. Online differentiation of mental arithmetic and rest | | MC | | fNIRS |
| Arsalidou and Taylor (2011) | Is 2+2=4? Meta-analyses of brain areas needed for numbers and calculations | | MC | | FMRI [meta analysis] |
| Hamada et al. (2018) | Comparison of brain activity between motor imagery and mental rotation of the hand tasks: a functional magnetic resonance imaging study | | | MR | fMRI |
| Kawamichi et al. (2007) | Distinct neural correlates underlying two- and three-dimensional mental rotations using three-dimensional objects | | | MR | fMRI |
| Shimoda et al. (2008) | Cerebral laterality difference in handedness: A mental rotation study with NIRS | | | MR | fNIRS |
| Harris and Miniussi (2003) | Parietal lobe contribution to mental rotation demonstrated with rTMS | | | MR | TMS |
| Khan and Hong (2017) | Hybrid EEG-fNIRS-Based Eight-Command Decoding for BCI: Application to Quadrucopter Control | WG | MC | MR | fNIRS and EEG |
| Tomasino and Gremese (2016) | Effects of Stimulus Type and Strategy on Mental Rotation Network: An Activation Likelihood Estimation Meta-Analysis | | | MR | PET + FMRI [meta analysis] |
| Herff et al. (2013) | Classification of mental tasks in the prefrontal cortex using fNIRS | WG | MC | MR | fNIRS |
| Khalaf et al. (2018) | Towards optimal visual presentation design for hybrid EEG—fTCD brain–computer interfaces | WG | | MR | EEG + fTCD |
| Qureshi et al. (2017) | Enhancing Classification Performance of Functional Near-Infrared Spectroscopy- Brain–Computer Interface Using Adaptive Estimation of General Linear Model Coefficients | | | MR | fNIRS |
| Hwang et al. (2014) | Evaluation of various mental task combinations for near-infrared spectroscopy-based brain-computer interfaces | | MC | MR | fNIRS |
| Roberts and Bell (2003) | Two- and three-dimensional mental rotation tasks lead to different parietal laterality for men and women | | | MR | EEG |
| Tagaris et al. (1998) | Functional magnetic resonance imaging of mental rotation and memory scanning: a multidimensional scaling analysis of brain activation patterns | | | MR | fMRI |
| Alivisatos and Petrides (1997) | Functional activation of the human brain during mental rotation | | | MR | PET |
| Friedrich et al. (2013) | Whatever Works: A Systematic User-Centered Training Protocol to Optimize Brain-Computer Interfacing Individually | WA | MC | MR | EEG |

*** WG: word generation; WA: word association*

*Table S4. Selected regions of interest for the LIT-based approach.*

| Inner Speech (IS) | Mental Calculation (MC) | Mental Rotation (MR) |
|---|---|---|
| *L \| Inferior Frontal Gyrus (p. Opercularis)* | *L-R \| Middle Frontal Gyrus* | *L-R \| Superior Parietal Lobule* |
| *L \| Inferior Frontal Gyrus (p. Triangularis)* | *R \| Angular Gyrus* | *L-R \| Inferior Parietal Lobule* |
| *L \| Superior Temporal Gyrus* | *L \| Superior Frontal Gyrus* | *L \| Precentral Gyrus* |
| *L \| Supramarginal Gyrus* | | *L-R \| Middle Frontal Gyrus* |
| *L \| Rolandic Operculum* | | *L-R \| Middle Occipital Gyrus* |
| *L \| Precentral Gyrus* | | |

*** L = left hemisphere; R = right hemisphere*

#### Mental-imagery task-pair selection process for fNIRS session

We carried out the task-pair selection at the individual subject level. For that, we first calculated the number of overlapping channels across all four layouts for each mental-imagery task, and selected the two tasks with the least number of overlapping channels. We computed the center of gravity (COG) for all four layouts per mental-imagery task in case this approach was not sufficient to select the two tasks (indicated with ? in Table S5). The mental tasks with the least number of overlapping channels and highest distance between them were the selected tasks.

***Table S5. Summary of the steps involved in selected the task-pair for each participant.***

| | *1st criterion: min # of overlapping channels between tasks* | | | Result → | | *2nd criterion: max distance between tasks* | | | Result → | | *Change selected task pair if original combination proofs incompatible* | | |
|---|---|---|---|---|---|---|---|---|---|---|---|---|---|
| | **Overlapping channels** | | | Selected task pair | | **Distance between COGs (mm)** | | | Selected task pair | | Conflict? | Selected task pair | |
| | IS | MC | MR | | | **IS** | **MC** | **MR** | | | | | |
| **P01** | 2 | 2 | 0 | ? | MR | 27,50 | 45,50 | 137,35 | MC | MR | No | MC | MR |
| **P02** | 0 | 0 | 2 | IS | MC | 136,18 | 173,11 | 106,48 | IS | MC | No | IS | MC |
| **P03** | 1 | 2 | 1 | IS | MR | 68,60 | 157,62 | 193,71 | IS | MR | No | IS | MR |
| **P04** | 2 | 1 | 0 | MC | MR | 107,60 | 187,46 | 195,29 | MC | MR | No | MC | MR |
| **P05** | 1 | 2 | 0 | IS | MR | 132,13 | 59,89 | 87,30 | IS | MR | **YES** | IS | MC |
| **P06** | 2 | 2 | 0 | ? | MR | 71,25 | 120,98 | 206,58 | MC | MR | No | MC | MR |
| **P09** | 2 | 2 | 0 | ? | MR | 143,84 | 152,83 | 212,09 | MC | MR | No | MC | MR |
| **P10** | 2 | 2 | 2 | ? | ? | 81,36 | 192,21 | 175,89 | MC | MR | No | MC | MR |
| **P11** | 3 | 2 | 2 | MC | MR | 75,69 | 74,12 | 123,46 | MC | MR | No | MC | MR |
| **P14** | 2 | 2 | 0 | ? | MR | 65,48 | 145,50 | 226,89 | MC | MR | No | MC | MR |
| **P15** | 2 | 2 | 2 | ? | ? | 72,95 | 155,34 | 154,82 | MC | MR | No | MC | MR |
| **P16** | 1 | 1 | 2 | IS | MC | 107,86 | 220,43 | 164,65 | IS | MC | **YES** | MC | MR |
| **P17** | 0 | 2 | 0 | IS | MR | 112,77 | 163,57 | 237,51 | IS | MR | **YES** | MC | MR |
| **P19** | 1 | 2 | 2 | IS | ? | 213,88 | 147,34 | 150,48 | IS | MR | **YES** | MC | MR |
| **P20** | 3 | 2 | 0 | MC | MR | 48,38 | 138,12 | 189,18 | MC | MR | No | MC | MR |
| **P21** | 2 | 2 | 1 | ? | MR | 201,47 | 139,88 | 207,34 | IS | MR | No | IS | MR |

Subject-specific optode layout for the selected mental-imagery task pair

Figure S6 illustrates schematically the selected optode layout for each participant (top view). The optode layouts for each mental-imagery task have been separated in these plots for clarity.

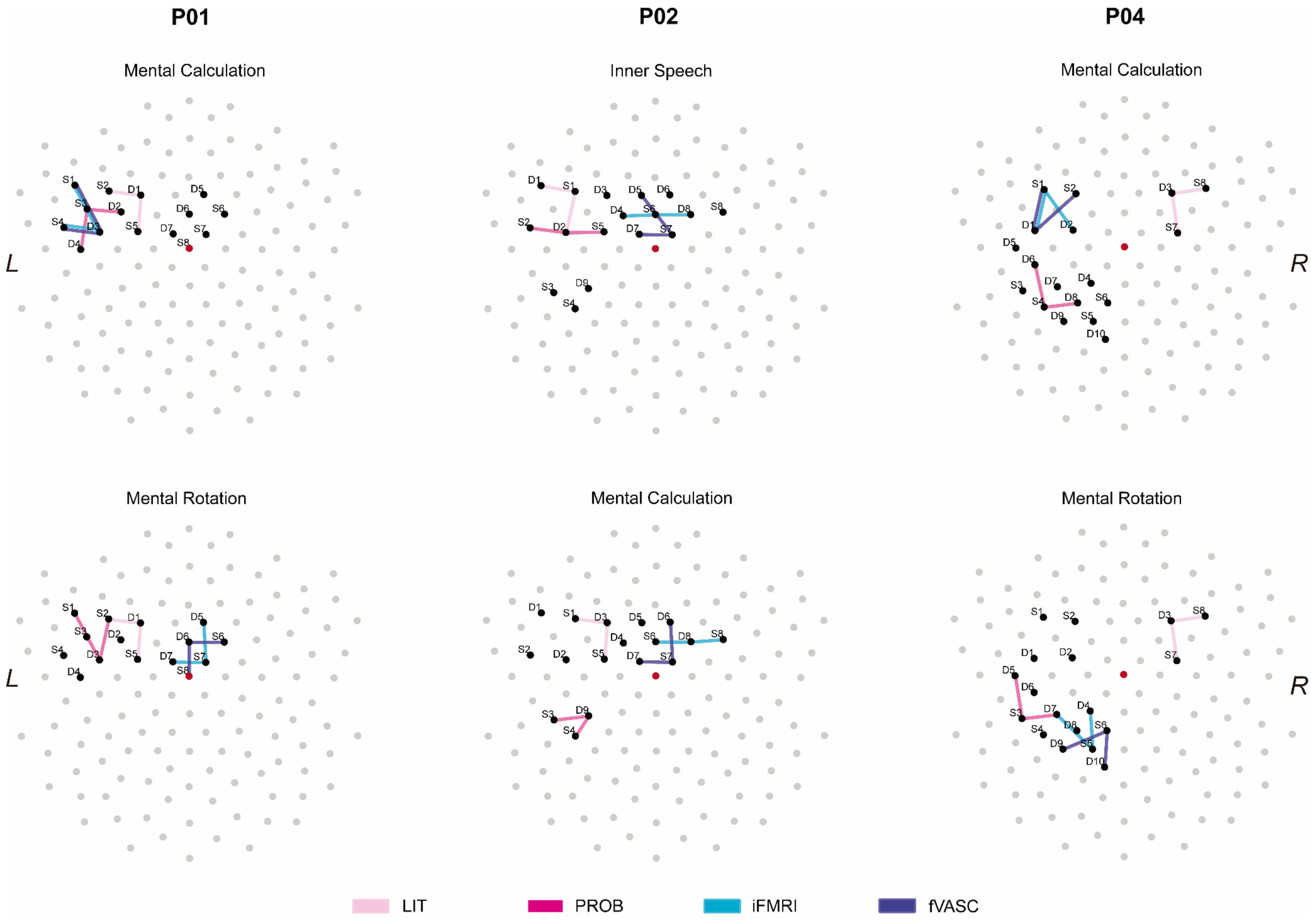
P01
P02
P04
Mental Calculation
Inner Speech
Mental Calculation
Mental Rotation
Mental Calculation
Mental Rotation
L
R
LIT
PROB
iFMRI
fVASC

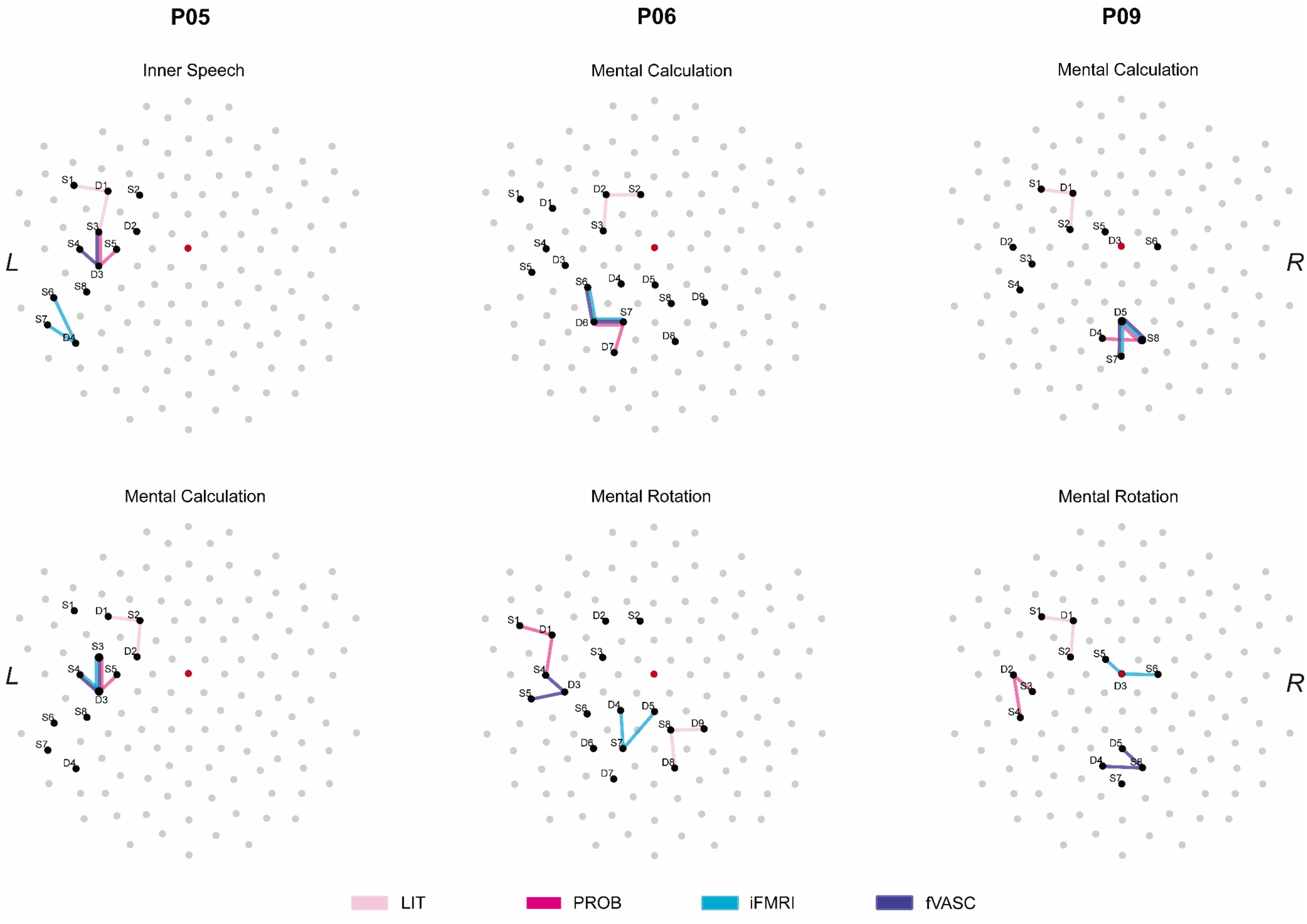
P05
P06
P09
Inner Speech
Mental Calculation
Mental Calculation
Mental Calculation
Mental Rotation
Mental Rotation
L
R
L
R
LIT
PROB
iFMRI
fVASC

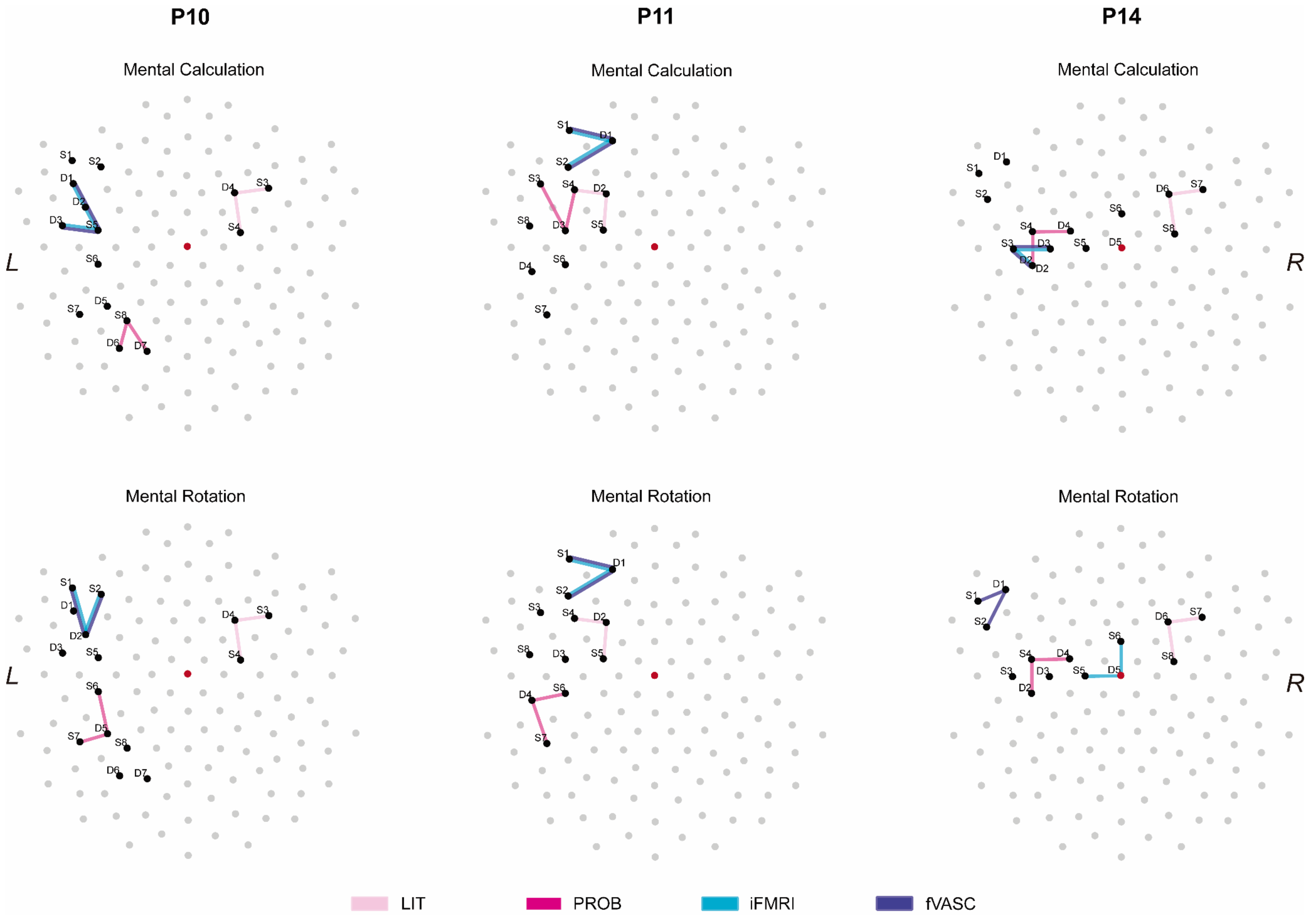
P10
P11
P14
Mental Calculation
Mental Rotation
L
R
LIT
PROB
iFMRI
fVASC

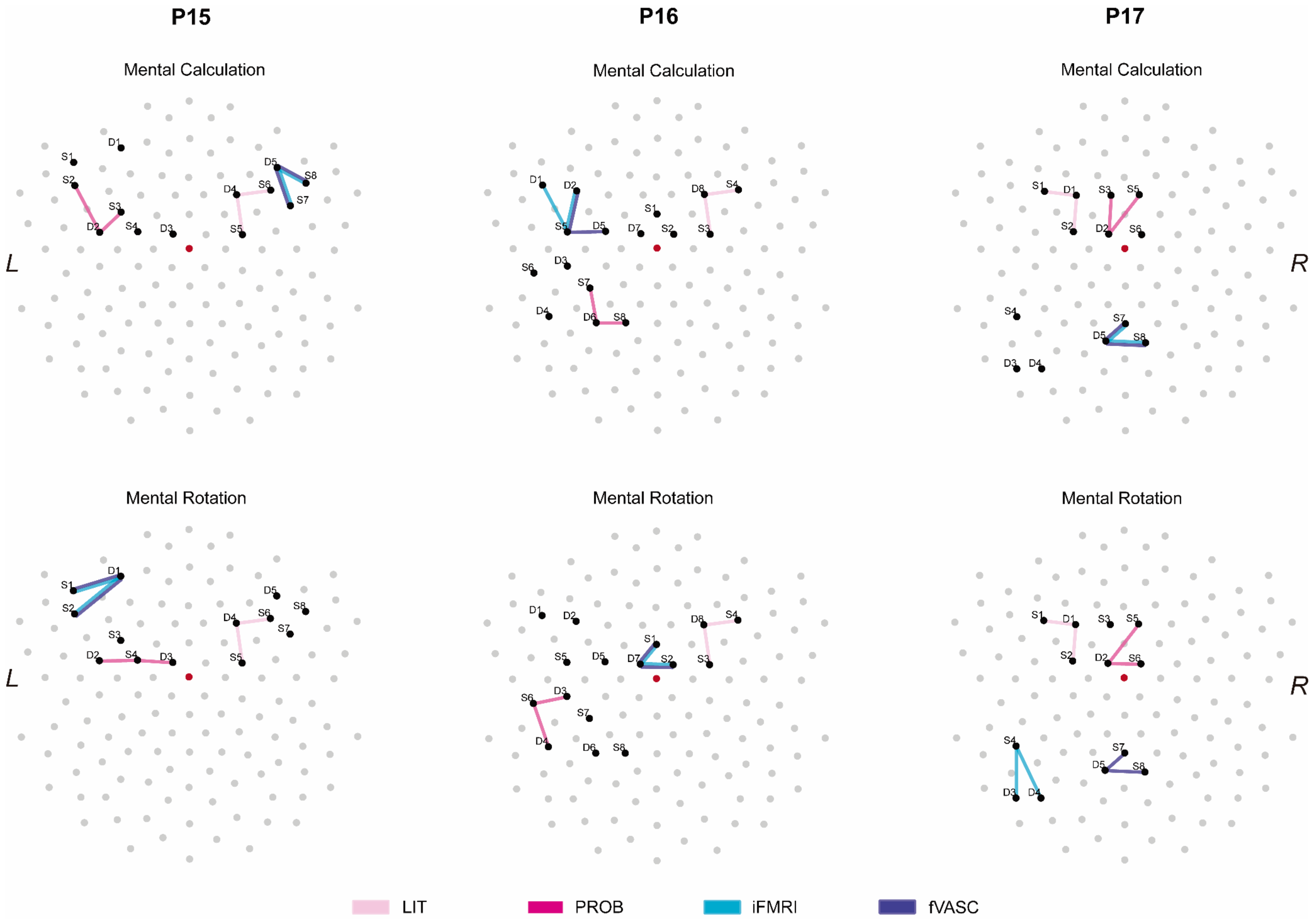
P15
P16
P17
Mental Calculation
Mental Calculation
Mental Calculation
Mental Rotation
Mental Rotation
Mental Rotation
L
R
L
R
LIT
PROB
iFMRI
fVASC

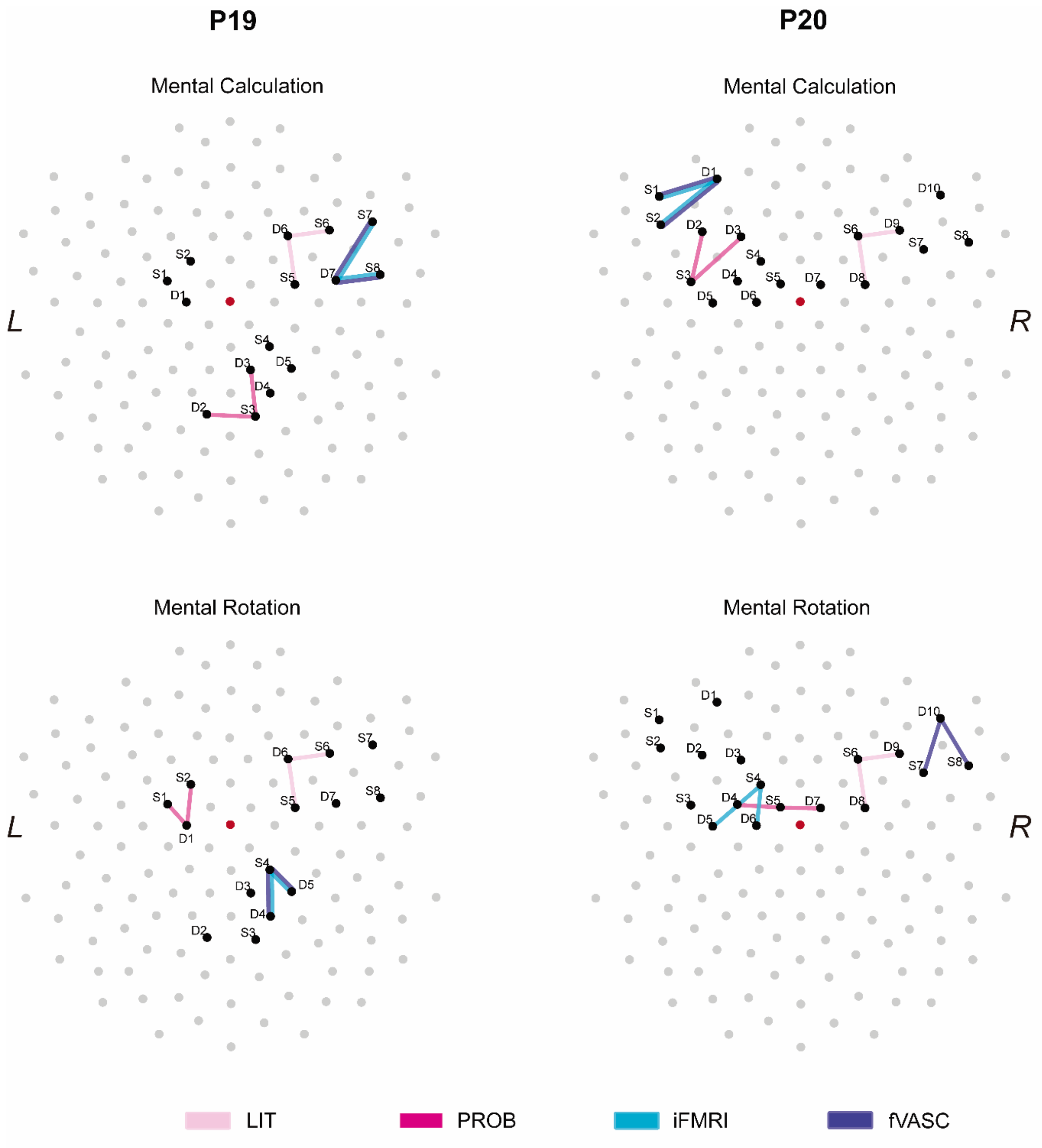


***Fig. S6. Subject- and approach-specific optode layouts for each mental-imagery task (top view).*** *Line colors represent the approach used to generate a given layout. 'S' represents sources and 'D' represents detectors. Cz location is marked in red.*

##### *A.3 Data analysis*

Data quality and presence of motion artifacts assessment

We computed the coefficient of variation (CV) to quantify the signal quality in each channel. Channels with CV >=7.5% were discarded from subsequent analyses. Figure S7 shows the percent of channels that fulfilled the CV criterion for each participant.

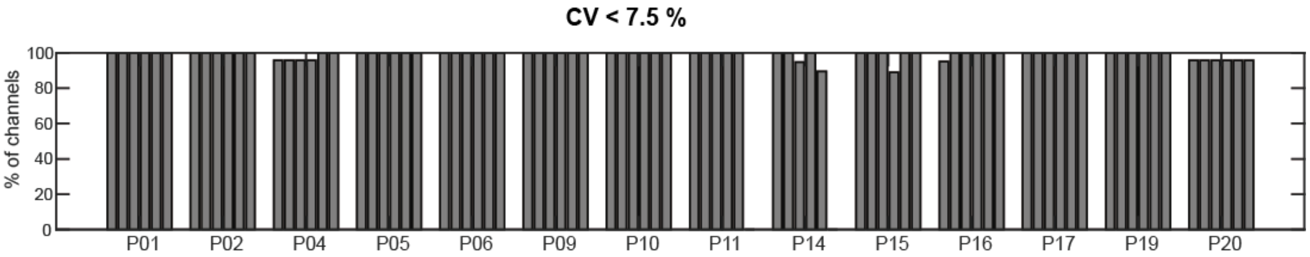


***Fig. S7. Signal quality****. This figure shows the percent of channels (y-axis) across runs and participants (x-axis) that survived the CV threshold of 7.5%. Almost all channels met the CV criterion across participants.*

The top panel of figure S8 summarizes the detected motion events per channel and run for each participant. The bottom panel provides cumulative motion events across channels and runs for participant P14.

#### B. Results

##### *B.1 Frequency maps*

We computed frequency maps for each mental-imagery task and approach to assess the spatial agreement of the selected channels across participants. The frequency maps shown in Figure S9 indicate that the selected channels varied considerably across subjects for PROB, iFMRI and fVASC approaches. In addition, iFMRI and fVASC approaches (the two most individualized ones) show the highest and most similar spatial extension for MC and MR. It is important to note that the (low) variability observed in the LIT approach is due to the use of the minimal individual anatomical information during the channel selection step (see Section 2.3.2.2).

##### *B.2 Examples of typical and weak/inverted hemodynamic responses*

Figure S10-A shows examples of four participants with typical hemodynamic responses (a positive deflection in HbO and a negative deflection in HbR) for a given approach, together with the projected activation on individual cortex reconstructions. Figure S10-B shows examples of four participants with weak/inverted hemodynamic responses.

#### C. Discussion

##### *C.1 Correspondence between fMRI and fNIRS block averages*

Figure S11 shows the correspondence between fMRI and fNIRS block averages calculated from channels placed according to the four approaches, for participants P01 and P16 during MR and MC tasks, respectively. We used channel-specific projection weights and projection spheres to compute spatially weighted fMRI block averages of the regions where the fNIRS signal most likely originated (see section 2.3.5.8 for details). The figure shows that channels placed based on approaches that use more individualized information show a better agreement with fMRI block averages. In addition, channels in close proximity can capture considerably different responses (see P01).

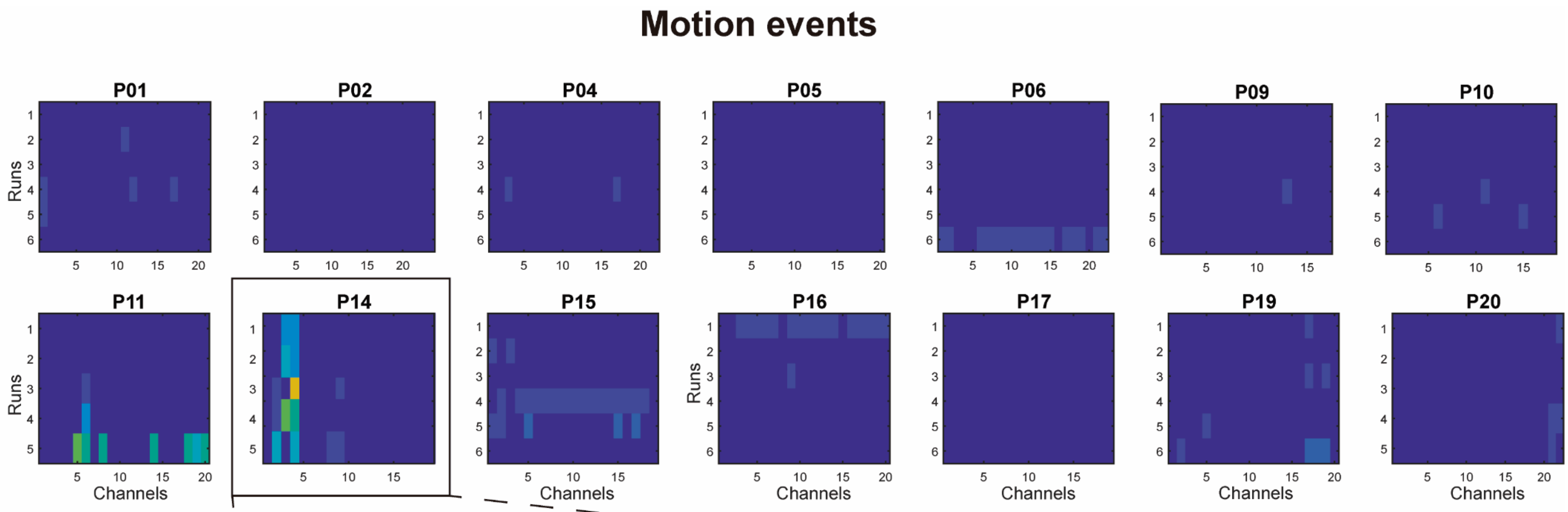


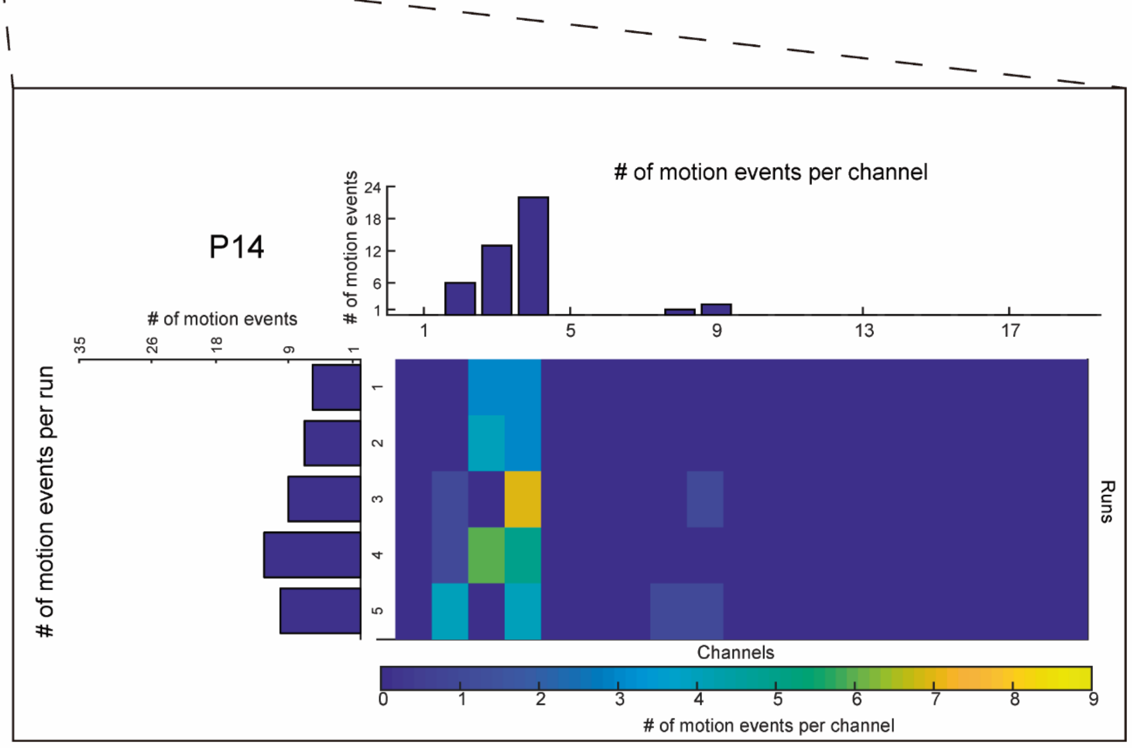


***Fig. S8. Motion artifacts*. *Top.*** *Presence of motion artifacts across channels (x-axis in subplot), runs (y-axis in subplot) across participants. The color in each cell indicates the frequency of detected motion artifacts within a run for a given channel. Note that the number of channels and runs were different across participants.* ***Bottom.*** *Closer look into the motion artifacts in participant P14. The top histogram show the cumulative number of motion events across runs for a given channel, while the histogram on the left shows the cumulative motion effects over channels for a given run.*

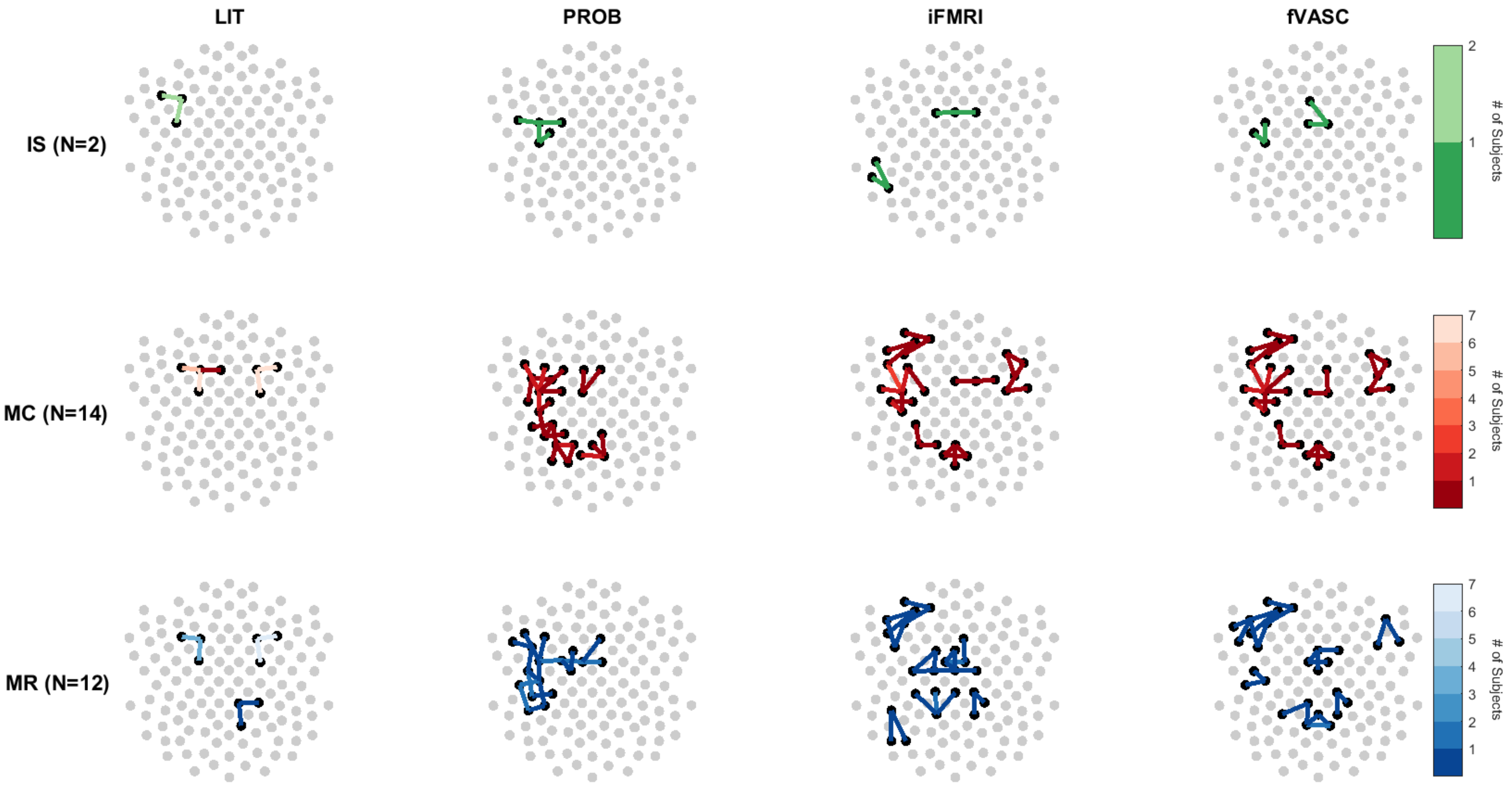


***Fig. S9. Top view of the channel frequency maps for each mental-imagery task (rows) and approach (columns).*** *Black dots indicate the locations where optodes were placed, while grey dots represent all 130 locations optodes could be located. Channels are indicated with lines and their colors indicate the number of participants who used a given channel. The selected channels vary considerably across subjects for PROB, iFMRI and fVASC approaches. iFMRI and fVASC approaches show highest similarity and spatial extent.*

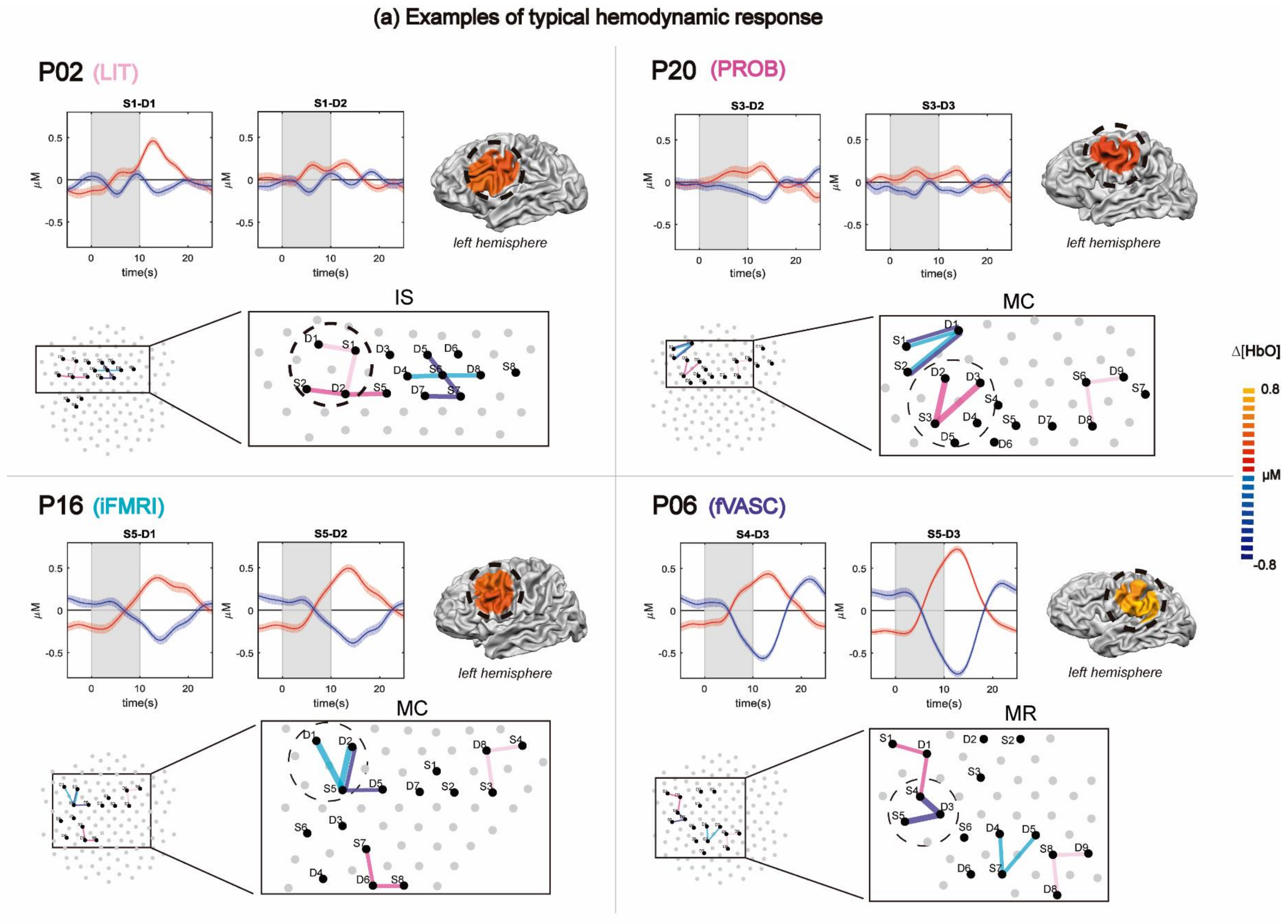
(a) Examples of typical hemodynamic response
P02 (LIT)
S1-D1
S1-D2
left hemisphere
IS
P20 (PROB)
S3-D2
S3-D3
left hemisphere
MC
Δ[HbO]
0.8
μM
-0.8
P16 (iFMRI)
S5-D1
S5-D2
left hemisphere
MC
P06 (fVASC)
S4-D3
S5-D3
left hemisphere
MR
time(s)
μM

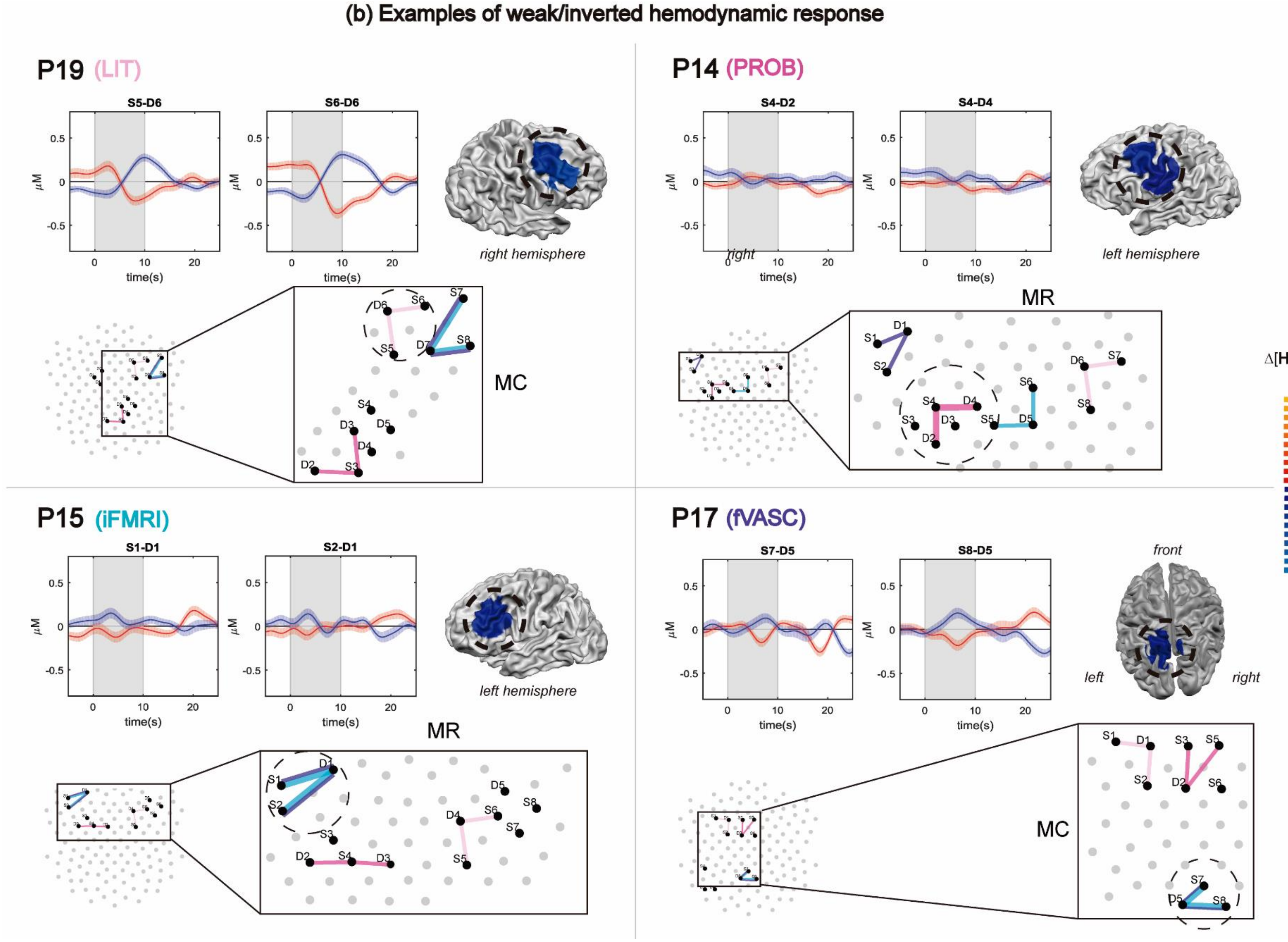


***Fig S10. Examples of typical (a) and weak/inverted (b) hemodynamic responses, for every approach-specific layout and mental-imagery tasks**. Every plot depicts a different participant. On the top part, fNIRS block averages of the two channels comprising a given layout are shown, with Δ[HbO]signal in red and Δ[HbR]signal in blue. The grey area indicates the onset and duration of the mental-imagery task (MC = Mental Calculation; MR= Mental Rotation). The bottom left plot shows the approach-specific optode layout for the same mental-imagery task as the block averages. The bottom right plot illustrates the projection of the two fNIRS channels in the individual anatomical data (3D surface reconstructions) for Δ[HbO].*

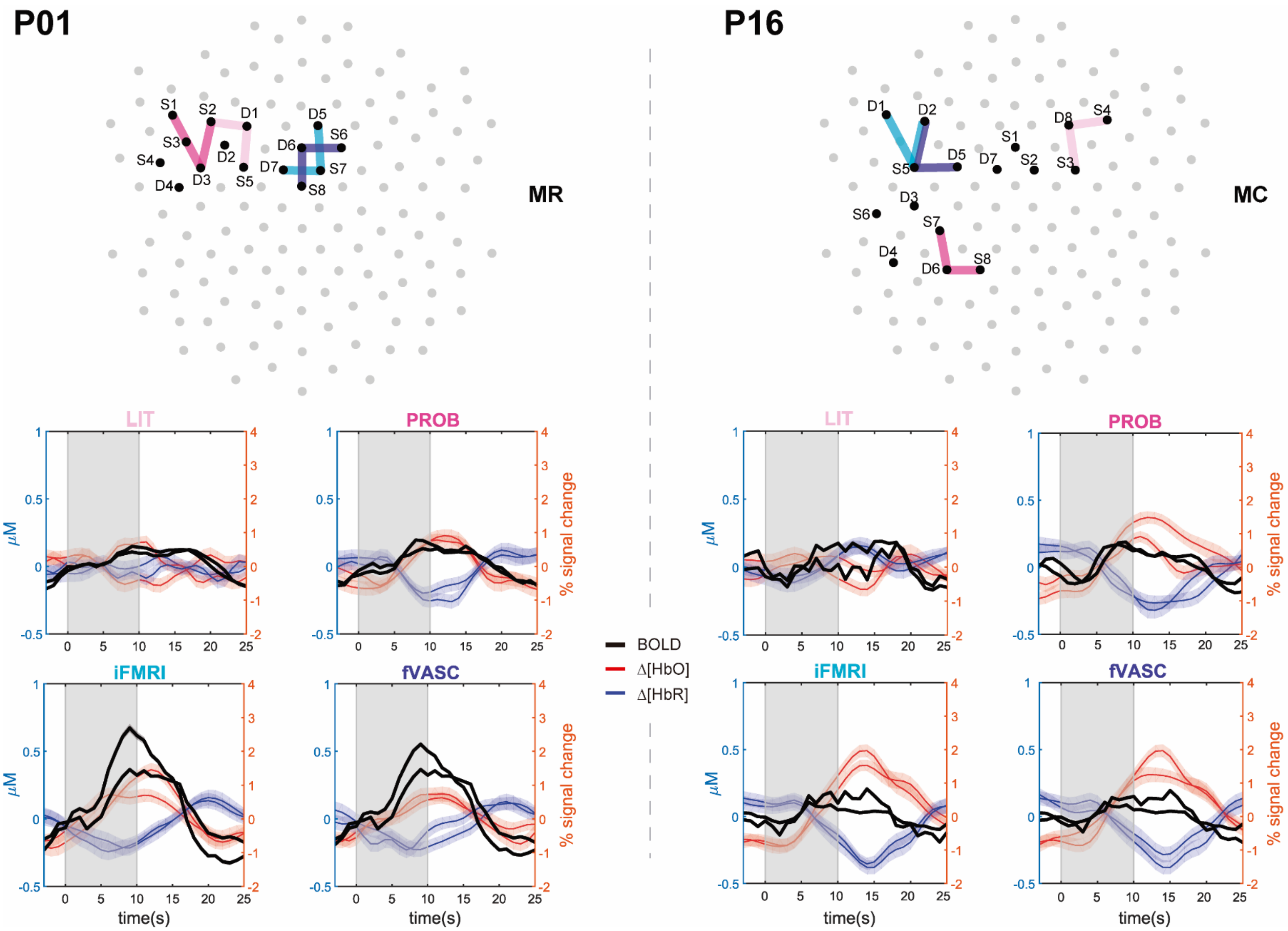


*__Fig. S11. Correspondence between fMRI and fNIRS block averages across different approaches in participants P01 (left) and P16 (right) for one of the mental-imagery task they performed (mental rotation [MR] and mental calculation [MC], respectively).__ The y-axis on the left represents concentration changes in μM for Δ[HbO] (red) and Δ[HbR] (blue) data, while the y-axis on the right represents the percent signal change for the fMRI time course (BOLD response, black line). The 0 value in the x-axis represents the task onset time (in s) and the gray area depicts the task duration. fNIRS time courses were normalized by the peak value before computing the block averages. The fNIRS block averages derived from the more individualized approaches show better agreement with fMRI data.*
